# Elicitation of broadly protective sarbecovirus immunity by receptor-binding domain nanoparticle vaccines

**DOI:** 10.1101/2021.03.15.435528

**Authors:** Alexandra C. Walls, Marcos C. Miranda, Minh N. Pham, Alexandra Schäfer, Allison Greaney, Prabhu S. Arunachalam, Mary-Jane Navarro, M. Alejandra Tortorici, Kenneth Rogers, Megan A. O’Connor, Lisa Shireff, Douglas E. Ferrell, Natalie Brunette, Elizabeth Kepl, John Bowen, Samantha K. Zepeda, Tyler Starr, Ching-Lin Hsieh, Brooke Fiala, Samuel Wrenn, Deleah Pettie, Claire Sydeman, Max Johnson, Alyssa Blackstone, Rashmi Ravichandran, Cassandra Ogohara, Lauren Carter, Sasha W. Tilles, Rino Rappuoli, Derek T. O’Hagan, Robbert Van Der Most, Wesley C. Van Voorhis, Jason S. McLellan, Harry Kleanthous, Timothy P. Sheahan, Deborah H. Fuller, Francois Villinger, Jesse Bloom, Bali Pulendran, Ralph Baric, Neil King, David Veesler

**Affiliations:** Department of Biochemistry, University of Washington, Seattle, WA 98195, USA; Institute for Protein Design, University of Washington, Seattle, WA 98195, USA; Department of Epidemiology, University of North Carolina at Chapel Hill, Chapel Hill, NC 27514, USA; Basic Sciences and Computational Biology, Fred Hutchinson Cancer Research Center, Seattle, WA 98109, USA; Department of Genome Sciences, University of Washington, Seattle, WA 98109, USA; Institute for Immunity, Transplantation and Infection, Stanford University School of Medicine, Stanford University, Stanford, CA, USA; Institut Pasteur and CNRS UMR 3569, Unité de Virologie Structurale, Paris, France; New Iberia Research Center and Department of Biology, University of Louisiana at Lafayette, New Iberia, LA, 70560 USA; Washington National Primate Research Center, Seattle, WA 98121, USA; Department of Microbiology, University of Washington, Seattle, WA 98195, USA; Department of Molecular Biosciences, The University of Texas at Austin, Austin, TX 78712, USA; Division of Allergy and Infectious Diseases, Department of Medicine, University of Washington School of Medicine, Seattle, WA 98195, USA; GSK, Siena, Italy; GSK, Rockville, MD, USA; GSK, Rixensart, Belgium; Bill & Melinda Gates Foundation, Seattle, WA 98109, USA

## Abstract

Understanding the ability of SARS-CoV-2 vaccine-elicited antibodies to neutralize and protect against emerging variants of concern and other sarbecoviruses is key for guiding vaccine development decisions and public health policies. We show that a clinical stage multivalent SARS-CoV-2 receptor-binding domain nanoparticle vaccine (SARS-CoV-2 RBD-NP) protects mice from SARS-CoV-2-induced disease after a single shot, indicating that the vaccine could allow dose-sparing. SARS-CoV-2 RBD-NP elicits high antibody titers in two non-human primate (NHP) models against multiple distinct RBD antigenic sites known to be recognized by neutralizing antibodies. We benchmarked NHP serum neutralizing activity elicited by RBD-NP against a lead prefusion-stabilized SARS-CoV-2 spike immunogen using a panel of single-residue spike mutants detected in clinical isolates as well as the B.1.1.7 and B.1.351 variants of concern. Polyclonal antibodies elicited by both vaccines are resilient to most RBD mutations tested, but the E484K substitution has similar negative consequences for neutralization, and exhibit modest but comparable neutralization breadth against distantly related sarbecoviruses. We demonstrate that mosaic and cocktail sarbecovirus RBD-NPs elicit broad sarbecovirus neutralizing activity, including against the SARS-CoV-2 B.1.351 variant, and protect mice against severe SARS-CoV challenge even in the absence of the SARS-CoV RBD in the vaccine. This study provides proof of principle that sarbecovirus RBD-NPs induce heterotypic protection and enables advancement of broadly protective sarbecovirus vaccines to the clinic.

## Introduction

The emergence of SARS-CoV-2 in late 2019 resulted in the COVID-19 pandemic that brought the world to a standstill ^1^. Moreover, the recurrent spillovers of coronaviruses in humans along with detection of SARS-CoV-2-, SARS-CoV- and MERS-CoV-related coronaviruses in bats, suggest that future zoonotic transmission events may continue to occur ^2–4^. SARS-CoV-2 infects host cells through attachment of the viral transmembrane spike (S) glycoprotein to angiotensin-converting enzyme 2 (ACE2), followed by fusion of the viral and host membranes ^1,5–12^. The SARS-CoV-2 S protein is the primary target of neutralizing antibodies (Abs), and the immunodominant receptor-binding domain (RBD) accounts for greater than 90% of the neutralizing activity in COVID-19 convalescent sera ^13,14^. Numerous monoclonal Abs (mAbs) recognizing distinct antigenic sites on the RBD were isolated and shown to neutralize viral entry and protect small animals and non-human primates (NHPs) from SARS-CoV-2 challenge ^13,15–22^. As a result, SARS-CoV-2 S is the focus of nucleic acid, vectored, and protein subunit vaccines currently being developed and deployed ^23–29^.

Worldwide sequencing of SARS-CoV-2 clinical isolates has led to the identification of numerous mutations in the >730,000 genome sequences available to date (https://www.gisaid.org/). The SARS-CoV-2 S D614G mutation has become globally dominant and is associated with enhanced viral transmission and replication but does not significantly affect Ab-mediated neutralization ^30–33^. Conversely, some mutations found in circulating SARS-CoV-2 isolates were shown to promote escape from mAbs and to reduce neutralization by immune sera ^34–37^. As a result, formulation of mAb cocktails neutralizing a broader spectrum of circulating SARS-CoV-2 variants emerged as a promising strategy to overcome this issue ^15,34,38,39^. The recent emergence of several variants with numerous S mutations is especially concerning, specifically the B.1.1.7, B1.351, and P.1 lineages that originated in the UK, South Africa, and Brazil, respectively ^40–42^. Some of these mutations lead to significant reductions in the neutralization potency of NTD- and RBD-specific mAbs, convalescent sera and Pfizer/BioNTech BNT162b2- or Moderna mRNA-1273-elicited sera ^43–46^.

We recently described a multivalent subunit vaccine displaying the SARS-CoV-2 RBD (RBD-NP) in a highly immunogenic array using a computationally designed self-assembling protein nanoparticle ^47,48^. Vaccination with RBD-NP resulted in 10-fold higher neutralizing Ab titers in mice than the prefusion-stabilized S-2P trimer (which is used in most current vaccines) despite a 5-fold lower dose and protected mice against mouse-adapted SARS-CoV-2 (SARS-CoV-2-MA) challenge ^47,49^. Furthermore, RBD-NP elicited robust neutralizing Ab and CD4 T cell responses in NHPs and conferred protection against SARS-CoV-2 infection in the nose, pharynges, and bronchioles ^50^. RBD-NP is currently being evaluated in two phase I/II clinical trials (NCT04742738 and NCT04750343).

Although the S fusion machinery (S_2_ subunit) has higher sequence conservation than the RBD ^5,51,52^, the breadth of neutralization and protection provided by RBD-based vaccines remains unknown. The isolation of RBD-specific cross-reactive mAbs neutralizing SARS-CoV-2 and SARS-CoV suggests that RBD-based vaccines could in principle elicit Abs that neutralize distantly related sarbecoviruses ^18,19,53^. RBD-based vaccines are also unaffected by S mutations outside of the RBD, especially in the highly variable N-terminal domain (NTD) ^37,54–59^. Here, we explored dose-sparing strategies for the RBD-NP vaccine and evaluated the impact of genetic diversity among SARS-CoV-2 clinical isolates and sarbecoviruses on vaccine-elicited Ab responses. We further designed mosaic and cocktail sarbecovirus RBD-NPs that elicit broad and protective Ab responses against heterologous sarbecovirus challenge, which could represent the next generation of pan-sarbecovirus vaccines.

### Dose-sparing RBD-NP vaccination protects mice against SARS-CoV-2 challenge

Considering the unprecedented need for rapid global distribution of SARS-CoV-2 vaccines, we set out to investigate the ability of RBD-NP to induce neutralizing and protective Ab titers at lower doses than previously tested: after either a single 1 µg immunization or two 0.1 µg immunizations (RBD antigen dose). BALB/c mice were immunized intramuscularly at week 0 with 0.1 µg (ten mice) or 1 µg (twenty mice) of AddaVax-adjuvanted SARS-CoV-2 RBD-NP. Three weeks later, the mice primed with 0.1 µg of vaccine and half of the mice primed with 1 µg of vaccine were boosted. Two weeks post-prime, serum binding and neutralizing Ab titers were comparable for the two vaccine doses (neutralization geometric mean titer (GMT) ∼3×10^2^) as measured using an MLV pseudotyped virus assay ^5,60^ (**Fig. 1a, Extended Data Fig. 1a**). Two weeks post-boost, serum neutralizing Ab titers were comparable for the 0.1 µg and 1 µg groups (GMT 0.9×10^4^ and 2×10^4^, respectively) (**Fig. 1b, Extended Data Fig. 1b**). Furthermore, we observed that the magnitude of binding and neutralizing Ab titers increased over time for the mice that received a single immunization (neutralization GMT 2×10^3^).

**Figure 1.**
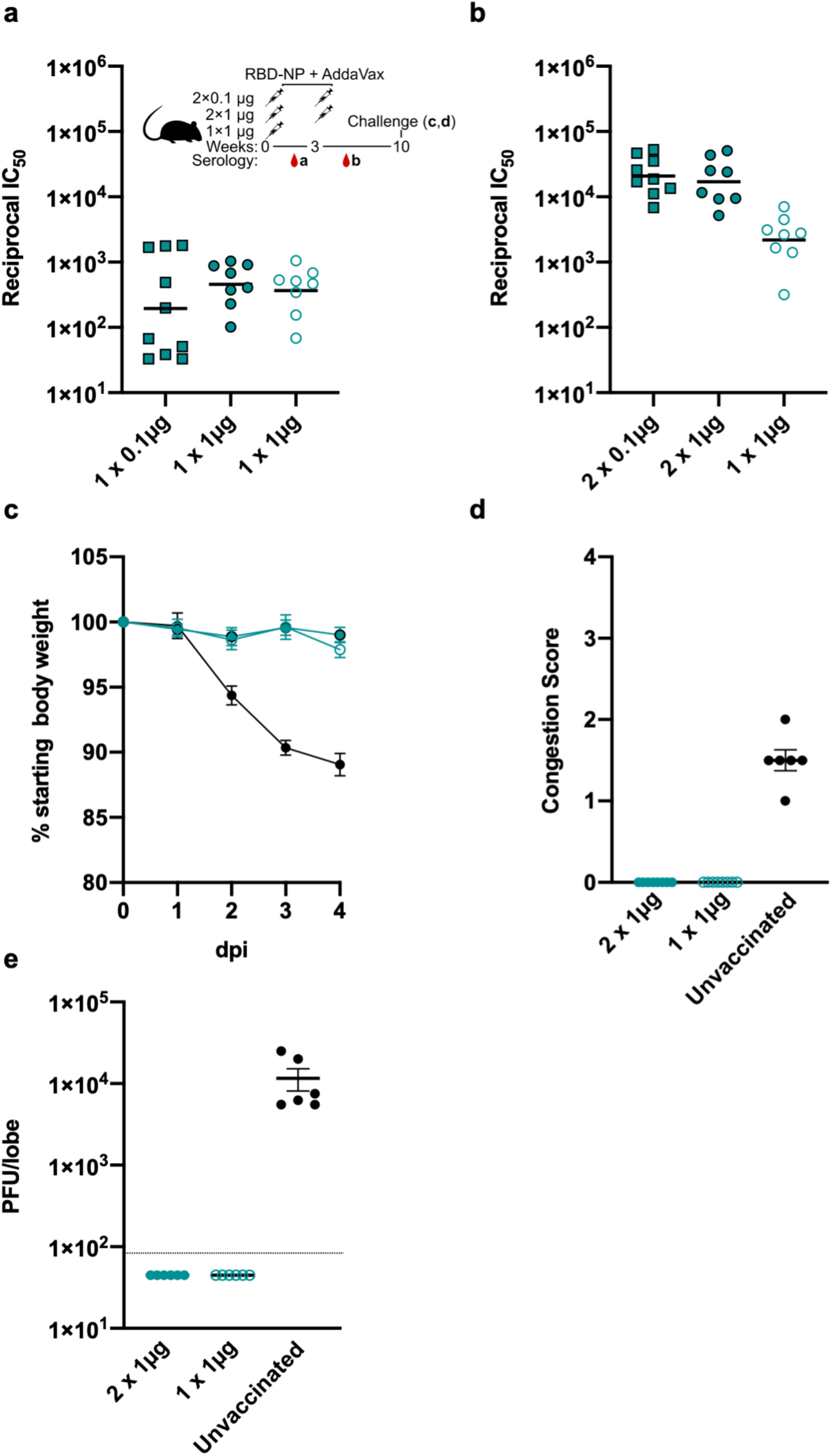
A single immunization with RBD-NP protects Balb/c cByJ mice from SARS-CoV-2 MA10 challenge. (a–b) Serum neutralizing Ab titers at 2 (a) or 5 (b) weeks post-prime determined using an MLV pseudotyping system; teal squares: 2×0.1 µg group; filled teal circles: 2×1 µg group; open teal circles: 1×1 µg group. (c) Weight loss following SARS-CoV-2 MA10 challenge up to 4 days post infection (N=6). Black, naïve mice; filled teal circles, 2×1 µg group; open teal circles, 1×1 µg group. (d) Congestion score following SARS-CoV-2 MA10 infection with a score of 0 indicating unchanged lung color and 4 indicating a darkened and diseased lung. (e) Viral titers in the mice lungs (expressed in plaque forming units per lobe) following challenge.

Six mice from each of the two groups that received 1 µg of vaccine along with six unvaccinated mice were challenged 10 weeks post-prime with 10^5^ plaque-forming units (pfu) of mouse-adapted SARS-CoV-2 MA10 ^61^ and followed for 4 days to assess protection from disease. All mice in the RBD-NP vaccinated groups were protected from body weight loss throughout the duration of the experiment, whereas control mice lost 10% of their weight on average (**Figure 1c)**. Analysis of viral titers in lung tissues and lung pathology indicated that the vaccinated mice did not appear affected by SARS-CoV-2 MA10 challenge whereas the control mice showed high viral load and lung discoloration 4 days post-infection (**Figure 1d–e)**. These results indicate that a single immunization with SARS-CoV-2 RBD-NP results in sufficiently high titers of neutralizing Abs to confer protection against SARS-CoV-2 MA10-induced disease. Furthermore, mice vaccinated twice with a ten-fold lower dose of antigen (0.1 µg) had higher serum neutralizing antibody titers than mice receiving the single 1 µg dose, suggesting multiple dose-sparing regimens that could help achieve global vaccination.

**Extended Data Fig. 1:**
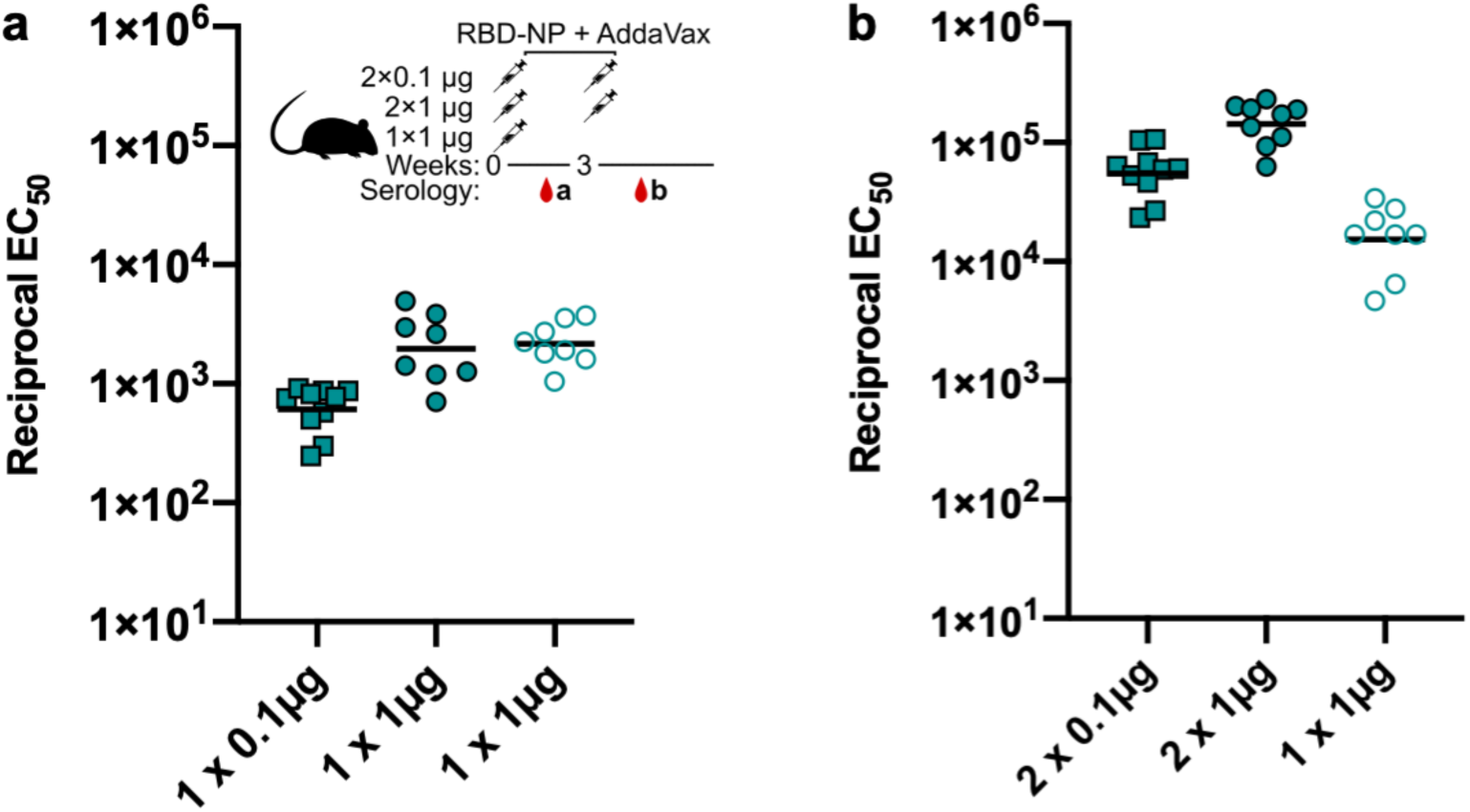
SARS-CoV-2 S 2P binding titers elicited by RBD-NP vaccination in mice. Antigen-specific Abs were measured 2 (a) or 5 (b) weeks post-prime; teal squares: 2×0.1 µg group; filled teal circles: 2×1 µg group; open teal circles: 1×1 µg group.

### SARS-CoV-2 RBD-NP vaccination induces Abs targeting diverse antigenic sites in NHPs

To characterize the epitopes targeted by vaccine-elicited polyclonal Abs, we used deep mutational scanning (DMS) with sera obtained 4 weeks post-boost from a pigtail macaque immunized at weeks 0 and 4 with a dose of AddaVax-adjuvanted RBD-NP containing 88 µg of the SARS-CoV-2 RBD (**Figure 2a**). These experiments rely on yeast surface display of RBD libraries covering nearly all possible amino acid mutations coupled with fluorescence-activated cell sorting to identify RBD mutants with attenuated Ab binding compared to the wild-type (Wuhan-1) SARS-CoV-2 RBD ^38,62^. No single mutation had more than a marginal effect on serum Ab (IgG/IgM/IgA) binding, indicating broad targeting of distinct RBD epitopes, whereas several COVID-19 human convalescent plasma analyzed by DMS displayed greater sensitivity to individual mutations ^14^ (**Figure 2b,c and Extended Data Fig. 2**). These results show that RBD-NP vaccination elicits diverse Ab responses that target multiple distinct antigenic sites and are more resilient to escape mutations than human convalescent plasma (HCP) ^13,14,21^.

**Figure 2:**
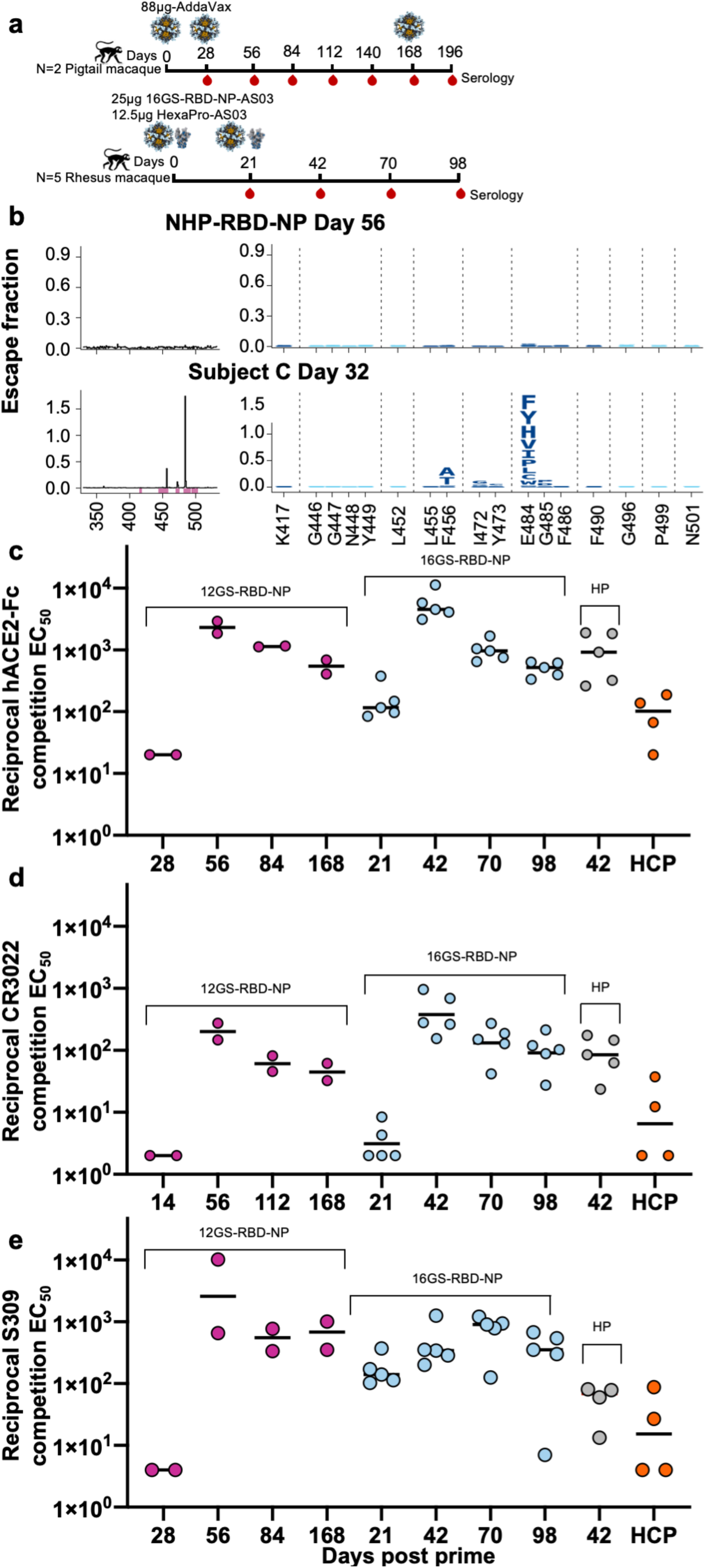
SARS-CoV-2 RBD-NP vaccination elicits high titers of Abs targeting diverse RBD antigenic sites in NHPs. (a) Vaccination schedules and RBD antigen doses of the different immunogens. (b) Effects of mutations on RBD binding by polyclonal serum Abs, measured by DMS analysis of serum obtained 8 weeks post-prime from a pigtail macaque vaccinated with 12GS-RBD-NP, compared to a previously reported DMS measurement on plasma Ab binding from a representative SARS-CoV-2 human convalescent individual reproduced here for comparison ^64^. The line plots on the left show the summed effect of all mutations at a site in the RBD on sera or plasma binding, with RBD site on the x axis and Ab escape on the y axis. Due to the use of sample-specific FACS gates, the y axes are scaled independently. Sites in the logo plots are colored dark blue if located in the receptor-binding ridge or cyan if located in the RBD 443–450 loop. Larger values indicate more Ab escape. (c–e) Competition ELISA between 0.2 nM human ACE2-Fc (c), 2 nM CR3022 mAb (d), or 0.01 nM S309 mAb (e) and RBD-NP-elicited sera in rhesus/pigtail macaques or HexaPro-elicited sera in rhesus macaques at various time points following vaccination, benchmarked against HCP. Each plot shows the magnitude of inhibition of ACE2/mAb binding to immobilized SARS-CoV-2 S 2P, expressed as reciprocal serum dilution blocking 50% of the maximum binding response.

To measure the magnitude of vaccine-elicited Abs against distinct RBD antigenic sites, we used quantitative competition ELISAs with two structurally characterized mAbs covering antigenic sites II (CR3022) and IV (S309) as well as human ACE2-Fc (hACE2-Fc), which binds to the receptor-binding motif (RBM), comprising antigenic sites Ia and Ib (**Figure 2d–f**). These experiments used sera from NHPs immunized twice with RBD-NP formulated with either AddaVax or AS03 at two different doses, sera from NHPs that received AS03-adjuvanted prefusion-stabilized HexaPro S trimer ^63^, and COVID-19 HCP. 3-4 weeks post-boost, all NHP sera had high titers of Abs targeting site Ia and Ib in the immunodominant RBM (site I, competition EC_50_s ≥ 1×10^3^), a correlate of neutralization potency ^13^ (**Figure 2d**), in agreement with the potent immunogenicity and protective efficacy of RBD-NP and HexaPro in NHPs ^50^. We also observed strong Ab responses against antigenic sites II and IV (competition EC_50_s ≥ 1×10^2^) (**Figure 2e–f**), which comprise conserved sarbecovirus epitopes recognized by neutralizing mAbs such as S2X35 (site II) and S309 (site IV) ^13,18^. Ab responses were durable against all three antigenic sites regardless of dose or adjuvant, with RBM-directed Ab titers decreasing by roughly half an order of magnitude 98–168 days post prime. RBD-NP elicited the highest peak binding titers towards all RBD antigenic sites evaluated compared to HexaPro and HCP, showcasing the potency and diversity of Ab responses induced by multivalent display of the RBD.

**Extended Data Fig. 2:**
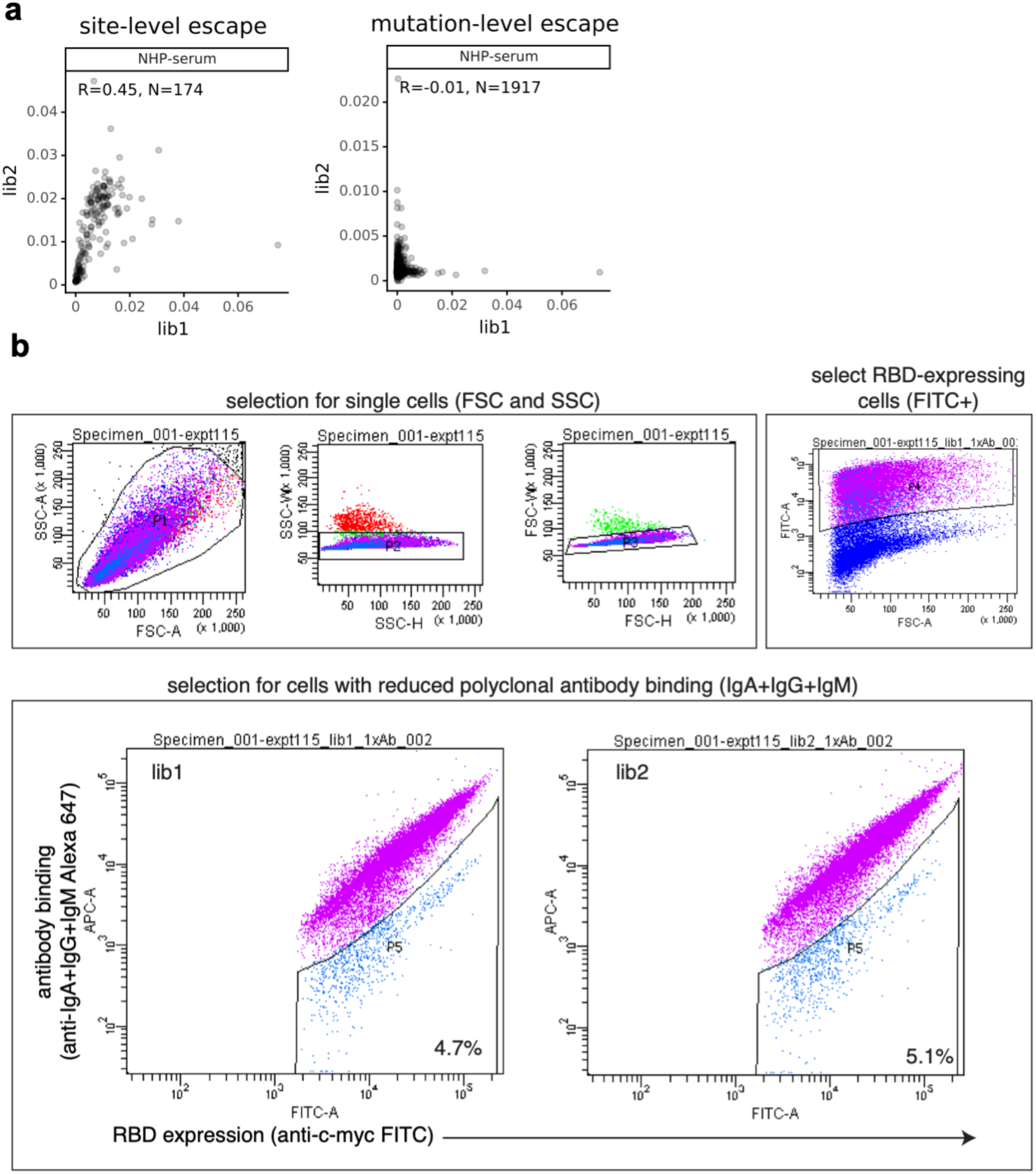
Effects of mutations on binding of HCP antibodies to RBD and FACS gating strategy. (a) Correlation plots of site- and mutation-level escape for each of the two independent RBD mutant libraries for the Ab-escape map shown in Fig. 2b. Site-level escape is the sum of the escape fractions for each mutation at a site. (b) Hierarchical FACS gating strategy used for selecting yeast cells expressing antibody-escape RBD variants. First, gates are selected to enrich for single cells (SSC-A vs. FSC-A, and FSC-W vs. FSC-H) that also express RBD (FITC-A vs. FSC-A, cells in pink). Second, cells expressing RBD mutants with reduced polyclonal Ab binding, detected with an anti-IgA+IgG+IgM secondary Ab, were selected with a gate that captured the ∼5% of cells with the lowest Ab binding (cells in blue).

### SARS-CoV-2 RBD-NP elicits potent neutralizing Ab responses in NHPs against a panel of variants

To assess the neutralization breadth of RBD-NP-elicited Abs, we evaluated serum neutralizing activity against a panel of pseudotyped viruses comprising wild-type (D614G) SARS-CoV-2 S and nine single-residue SARS-CoV-2 RBD mutants detected in clinical isolates (G446S, Y453F, L455F, T478I, E484A/K, F486L, S494P, and N501Y) as well as the B.1.1.7 (H69-V70 deletion, Y144 deletion, N501Y, A570D, P681H, T716I, S982A, D1118H) and B.1.351 (L18F, D80A, D215G, L242-L244 deletion, R246I, K417N, E484K, N501Y, A701V) variants of concern that originated in the UK and South Africa, respectively ^40,41^.

Most single-residue RBD mutations tested did not affect the serum neutralizing titers of NHPs vaccinated twice with RBD-NP or soluble HexaPro using an HIV pseudotyped virus compared to wild-type (D614G) SARS-CoV-2 S (GMT 5×10^2^ and 7×10^2^ for RBD-NP and HexaPro, respectively; **Fig. 3a**). The N501Y substitution present in the B.1.1.7 and B.1.351 lineages, the mink-associated Y453F substitution, and the prevalent N439K mutation did not affect the neutralization potency of any RBD-NP- or HexaPro-elicited sera significantly, while these substitutions have been associated with loss of neutralization for some mAbs ^44,46,65,66^. The E484K mutation reduced serum neutralizing activity by ∼5-fold and ∼7-fold for RBD-NP and HexaPro (GMT 1×10^2^) compared to wild-type (D614G) SARS-CoV-2 S, respectively, whereas the E484A and E484D mutations did not significantly affect the neutralizing activity induced by either immunogen. These experiments suggest that a substantial part of the neutralizing activity elicited by both RBD-NP and HexaPro is focused on the RBM, near position 484. To understand the impact of the full constellation of mutations present in S from the B.1.1.7 and B.1.351 lineages, we evaluated vaccine-elicited serum neutralizing activity against the corresponding HIV and VSV pseudotyped variant viruses. Although we did not observe any reduction in neutralization titers towards the B.1.1.7 variant, similar to what was seen with the N501Y point mutant, animals vaccinated with RBD-NP or HexaPro had 5-fold reduced serum neutralizing Ab titers against the B.1.351 S variant using HIV pseudovirus (GMT 1×10^2^; **Fig. 3b**) and nearly undetectable levels of neutralization using VSV pseudotyped viruses (**Fig. 3c**). These changes in neutralizing activity were presumably largely due to the E484K substitution albeit RBD-specific binding Ab titers were not markedly different compared to widtype RBD (**Extended Data Fig. 3a**). Furthermore, we observed a nearly ten-fold reduction in potency in sera from individuals who received two doses of the Pfizer-BioNTech BNT162b2 mRNA vaccine when comparing serum neutralization titers against B.1.351 S (GMT 6.7×10^1^) pseudotyped virus to wild-type (D614G) SARS-CoV-2 S (GMT 6×10^2^) (**Fig. 3b**). These findings show that, as is the case for many convalescent individuals ^13,64,67^, an important fraction of vaccine-elicited neutralizing Abs in NHPs and humans is focused on the RBM (specifically around position 484), independently of the immunogen (RBD-NP, HexaPro, or 2P-stabilized spike ^68^) or the vaccine modality (protein or mRNA). However, as neutralization assays may underestimate the contribution of NTD-specific neutralizing Abs ^54^, protection from challenge will be the ultimate readout.

**Figure 3.**
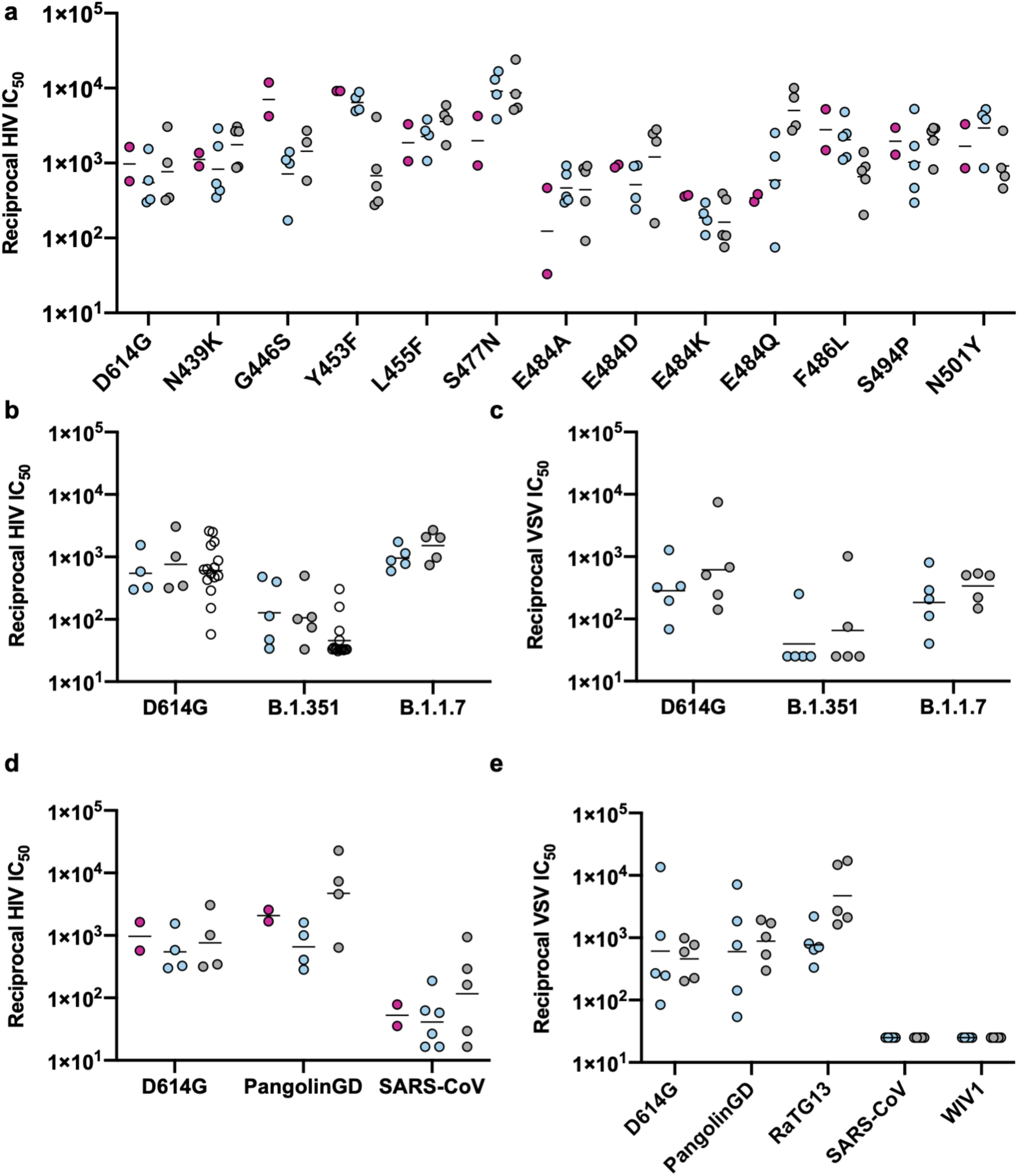
SARS-CoV-2 RBD-NP and HexaPro S elicit Abs with similar neutralization breadth towards SARS-CoV-2 RBD mutants detected in clinical isolates. (a) Neutralizing Ab titers against wildtype (D614G) SARS-CoV-2 S and RBD point mutants determined using SARS-CoV-2 RBD-NP-elicited sera in pigtail macaques (magenta), in rhesus macaques (blue), and HexaPro-elicited sera in rhesus macaques (gray) with an HIV pseudotyping system. (b) Neutralizing Ab titers against HIV pseudotyped viruses harboring wildtype (D614G) SARS-CoV-2 S, B.1.351 S or B.1.1.7 S determined using SARS-CoV-2 RBD-NP-elicited sera in rhesus macaques (blue), HexaPro-elicited sera in rhesus macaques (gray), or plasma from individuals who received two doses of Pfizer mRNA vaccine (open circles). (c) Neutralizing Ab titers against VSV pseudotyped viruses harboring wildtype (D614G) SARS-CoV-2 S, B.1.351 S or B.1.1.7 S determined using SARS-CoV-2 RBD-NP-elicited sera in rhesus macaques (blue) or HexaPro-elicited sera in rhesus macaques (gray). (d) Neutralizing Ab titers against HIV pseudotyped viruses harboring wildtype (D614G) SARS-CoV-2 S, Pangolin-GD S or SARS-CoV S determined using SARS-CoV-2 RBD-NP-elicited sera in pigtail macaques (magenta), in rhesus macaques (blue), or HexaPro-elicited sera in rhesus macaques (gray). (e) Neutralizing Ab titers against VSV pseudotyped viruses harboring wildtype (D614G) SARS-CoV-2 S, Pangolin-GD S, RaTG13 S, SARS-CoV S or WIV1 S determined using SARS-CoV-2 RBD-NP-elicited sera in rhesus macaques (blue) or HexaPro-elicited sera in rhesus macaques (gray).

To further investigate the relationships between neutralizing Ab titers and emerging SARS-CoV-2 variants, we vaccinated the two pigtail macaques a third time with SARS-CoV-2 RBD-NP, 24 weeks after the primary immunization. This boost induced very high serum neutralizing activity against wild-type (D614G) SARS-CoV-2 (GMT 2×10^5^) (**Extended Data Fig. 3b**) as well as the B.1.1.7 (GMT 2×10^4^) and B.1.351 (GMT 4×10^3^) variants of concern. These results are consistent with recent studies showing that boosting pre-existing immunity in COVID-19 convalescent individuals with a single mRNA vaccination elicited high (neutralizing) Ab titers, including against the B.1.351 variant ^69–71^, and suggest that a third vaccination of naïve individuals could be a suitable strategy to limit the impact of emerging variants while vaccines are updated.

### SARS-CoV-2 RBD-NP elicits cross-reactive sarbecovirus polyclonal Abs in NHPs

As RBD-NP-elicited polyclonal Abs proved resilient to a range of mutations observed in SARS-CoV-2 isolates, we next investigated cross-reactivity with a panel of sarbecovirus RBDs. SARS-CoV-2 RBD-NP-elicited polyclonal Abs (purified from pigtail macaque serum obtained 10 weeks post-prime) strongly cross-reacted with the SARS-CoV-2-related Pangolin-GD and RaTG13 RBDs and weakly bound the RmYN02, SARS-CoV, WIV16 and ZXC21 RBDs (**Extended Data Fig. 3c–d**). Measurement of Ab binding titers using the SARS-CoV and SARS-CoV-2 S 2P ectodomain trimers by ELISA showed that the SARS-CoV-2 RBD-NP and HexaPro induced similar levels of Abs against each antigen, with responses roughly two orders of magnitude higher against SARS-CoV-2 S 2P compared to SARS-CoV S 2P (**Extended Data Fig. 3e**). The two immunogens also elicited similar peak levels of ACE2-competing Abs, which correlate with neutralizing activity ^13^, in a competition ELISA assay using SARS-CoV S 2P as antigen (**Extended Data Fig. 3f**).

Motivated by the cross-reactivity of RBD-NP-elicited polyclonal Abs with sarbecovirus RBDs and the correlation between ACE2 competition and serum neutralization titers ^13^, we evaluated neutralization of a panel of HIV and VSV pseudoviruses harboring distinct sarbecovirus S glycoproteins. RBD-NP-elicited sera efficiently neutralized pseudotyped viruses harboring the S glycoprotein of the Pangolin-GD isolate (GMT HIV and VSV 6×10^2^) ^72^ (**Figure 3d–e)**, in agreement with the close phylogenetic relationship of its RBD with that of SARS-CoV-2 S. HexaPro-elicited NHP sera also inhibited Pangolin-GD (GMT HIV 7×10^2^ and VSV 6×10^2^) and RaTG13 pseudotypes (GMT VSV 8×10^2^) with slightly although not significantly higher potency than RBD-NP sera (**Figure 3d–e)**. Furthermore, we observed that both RBD-NP and HexaPro induced polyclonal Abs that weakly neutralized HIV pseudovirus carrying SARS-CoV S from the Urbani isolate (RBD-NP GMT 5×10^1^ and HexaPro GMT 1×10^2^) (**Figure 3d–e)**. Together, these data demonstrate that both immunogens elicited comparable breadth and potency against the sarbecoviruses tested.

**Extended Data Fig. 3.**
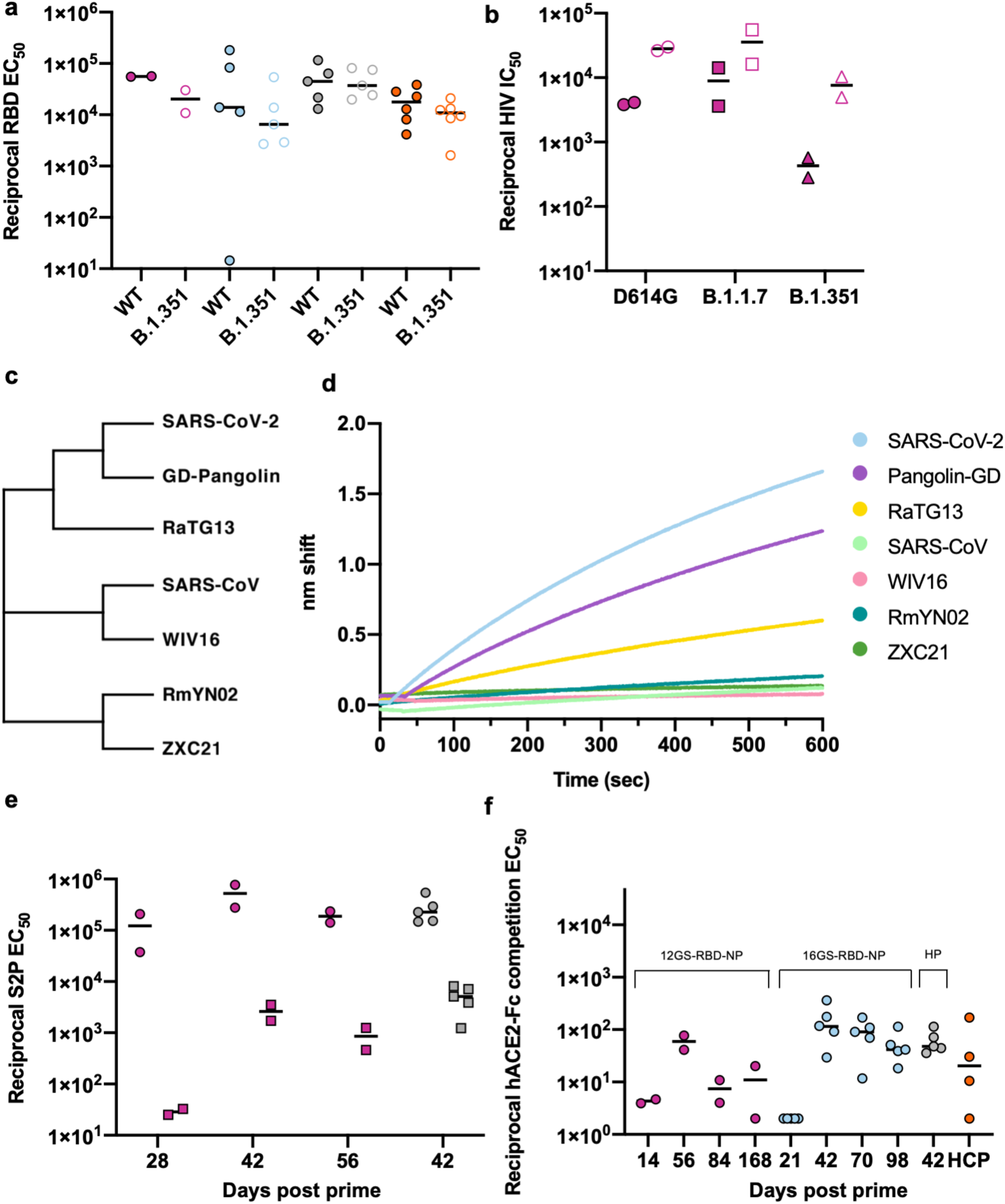
Evaluation of vaccine-elicited binding and neutralizing Ab titers against SARS-CoV-2 variants and distinct sarbecoviruses. (a) Wildtype (closed circles) and B.1.351 (open circles) SARS-CoV-2 RBD-specific Ab binding titers of SARS-CoV-2 RBD-NP-elicited sera in pigtail macaques (magenta) and in rhesus macaques (blue), HexaPro-elicited sera (gray), or HCP (orange) analyzed by ELISA. (b) Neutralizing Ab titers against wildtype (D614G) SARS-CoV-2 S, B.1.1.7, and B.1.351 S HIV pseudoviruses from NHPs immunized with 12GS-RBD-NP at day 56 (two immunizations, filled symbols) and 196 (three immunizations, open symbols) (c) Cladogram based on sarbecovirus RBD amino acid sequences. (d) Biolayer interferometry analysis of binding of 1 µM purified polyclonal pigtail macaque IgGs (obtained 70 days post prime) to sarbecovirus RBDs immobilized at the surface of biosensors. (e) SARS-CoV-2 S 2P (circles) or SARS-CoV S 2P (squares) Ab binding titers of SARS-CoV-2 RBD-NP-elicited sera in Pigtail macaques (magenta) or HexaPro-elicited sera in Rhesus macaques (gray) analyzed by ELISA. (f) Competition ELISA between 0.13 nM human ACE2-Fc and RBD-NP-elicited sera in pigtail macaques, rhesus macaques, or HexaPro S-elicited sera in rhesus macaques against SARS-CoV S 2P at various time points following vaccination benchmarked against COVID-19 HCP. Each plot shows the magnitude of inhibition of ACE2 binding to immobilized SARS-CoV S 2P, expressed as reciprocal serum dilution blocking 50% of the maximum binding response.

### Design, assembly and characterization of mosaic and cocktail sarbecovirus RBD-NPs

We and others have recently evaluated nanoparticle immunogens that display multiple antigenic variants of viral glycoproteins as a potential route to broadly protective vaccines ^73–76^. Given the large number of coronaviruses circulating in zoonotic reservoirs, such vaccines could be important for preventing future pandemics ^3,4^. We expressed and purified four proteins in which the RBDs from the S proteins of SARS-CoV-2, SARS-CoV, and the bat coronaviruses WIV1 and RaTG13 were genetically fused to the I53-50A trimer. An equimolar mixture of these four proteins was added to the I53-50B pentamer to assemble a mosaic RBD-NP (mRBD-NP) co-displaying the four RBDs on the same nanoparticle (**Fig. 4a** and **Extended Data Fig. 4a**). We also assembled a trivalent mosaic RBD-NP lacking the SARS-CoV RBD (mRBD-NP-DO), as well as cocktail immunogens in which three (cRBD-NP-DO; lacking the SARS-CoV RBD) or four (cRBD-NP) independently assembled nanoparticles, each displaying a single type of the aforementioned RBDs, were mixed after assembly. Finally, we made a bivalent mosaic RBD-NP co-displaying the SARS-CoV and SARS-CoV-2 RBDs (**Extended Data Fig. 4a**) to directly confirm co-display using a sandwich binding assay. All of the nanoparticle immunogens formed the intended icosahedral architecture and retained native antigenicity, as shown by SDS-PAGE, dynamic light scattering, electron microscopy analysis of negatively stained samples and binding to human ACE2 (**Extended Data Fig. 4b–e**). We found that the vaccine candidates were stable for at least four weeks at several temperatures except the highest temperature evaluated (37°C), at which we observed a decrease in ACE2 recognition over time, presumably due to aggregation (**Extended Data Fig. 5**). Following immobilization using the SARS-CoV-2-specific mAb S2H14, the quadrivalent, trivalent, and bivalent mRBD-NPs all bound the Fab of the SARS-CoV-specific Ab S230 ^13,77–79^, confirming co-display, whereas the monovalent SARS-CoV-2 RBD-NP did not (**Extended Data Fig. 6a**). We determined that the reactivity of the trivalent mRBD-NP derived from the inclusion of the WIV1 RBD in this vaccine, as the monovalent WIV1 RBD-NP also bound the S230 Fab after immobilization with hACE2 (as expected ^3^), whereas the monovalent SARS-CoV-2 and RaTG13 RBD-NPs did not (**Extended Data Fig. 6b**).

**Figure 4.**
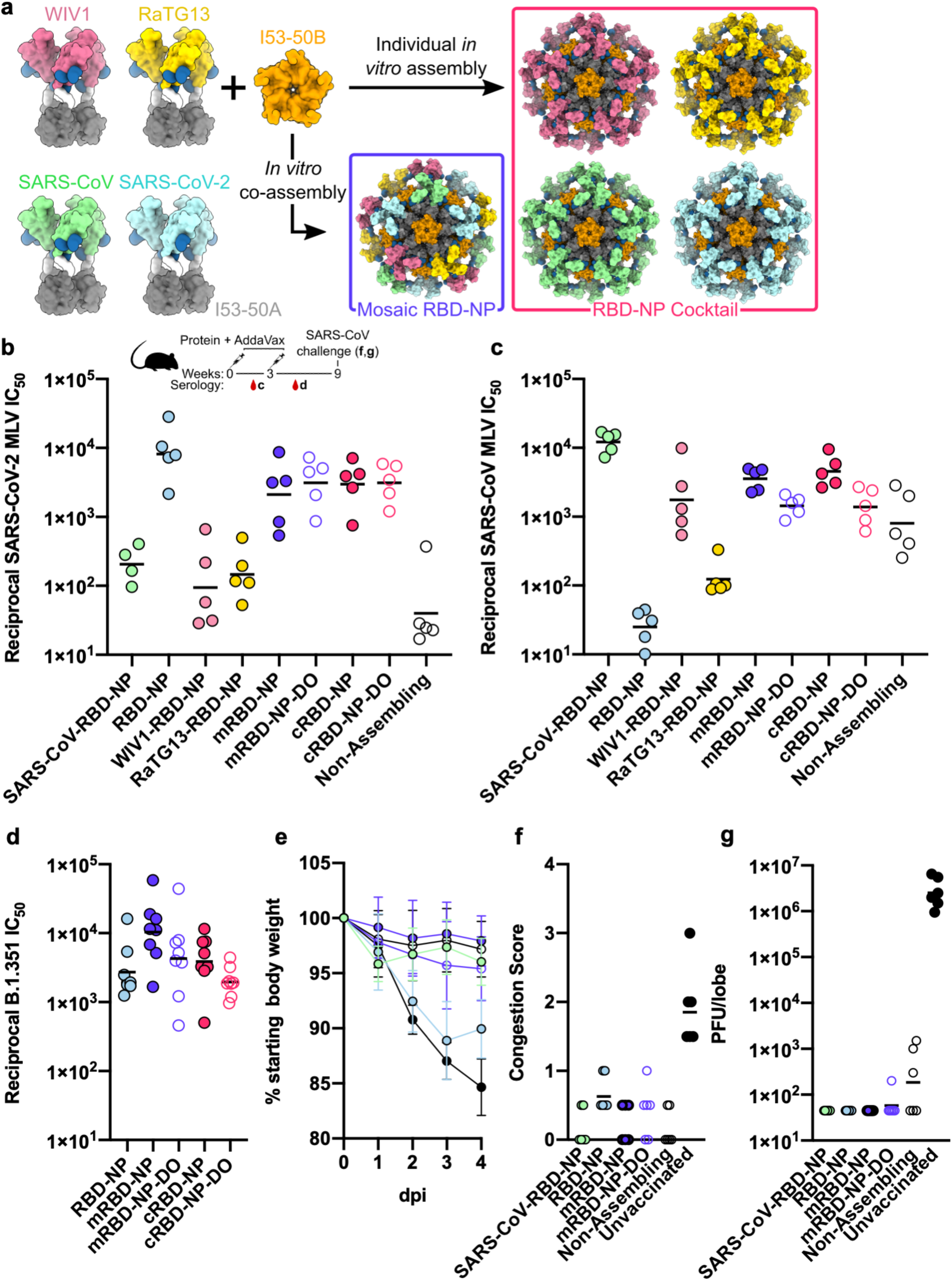
Mosaic and cocktail RBD-NPs elicit neutralizing Ab against multiple sarbecoviruses and protect against heterotypic challenge in 15 week old Balb/c cByJ mice. **a**, Schematic of *in vitr*o assembly of mRBD-NP and cRBD-NP. **b**, Neutralizing Ab titers against wildtype (D614G) SARS-CoV-2 S MLV pseudovirus at week 5 (2 weeks post-boost) elicited by monovalent, mosaic, and cocktail RBD-NPs. **c,** Neutralizing Ab titers against SARS-CoV S MLV pseudovirus at week 5 (2 weeks post-boost) elicited by monovalent, mosaic, and cocktail RBD-NPs. **d**, Neutralizing Ab titers against SARS-CoV-2 B.1.351 S MLV pseudovirus at week 5 (2 weeks post-boost) elicited by monovalent, mosaic, and cocktail RBD-NPs. **e**, Weight loss following SARS-CoV MA15 challenge up to 4 days post infection (N=6). **f**, Congestion score following SARS-CoV MA15 infection with a score of 0 indicating unchanged lung color and 4 indicating a darkened and diseased lung. **g,** Viral titers in the mice lungs (expressed in plaque forming units per lobe) following challenge.

**Extended Data Fig. 4:**
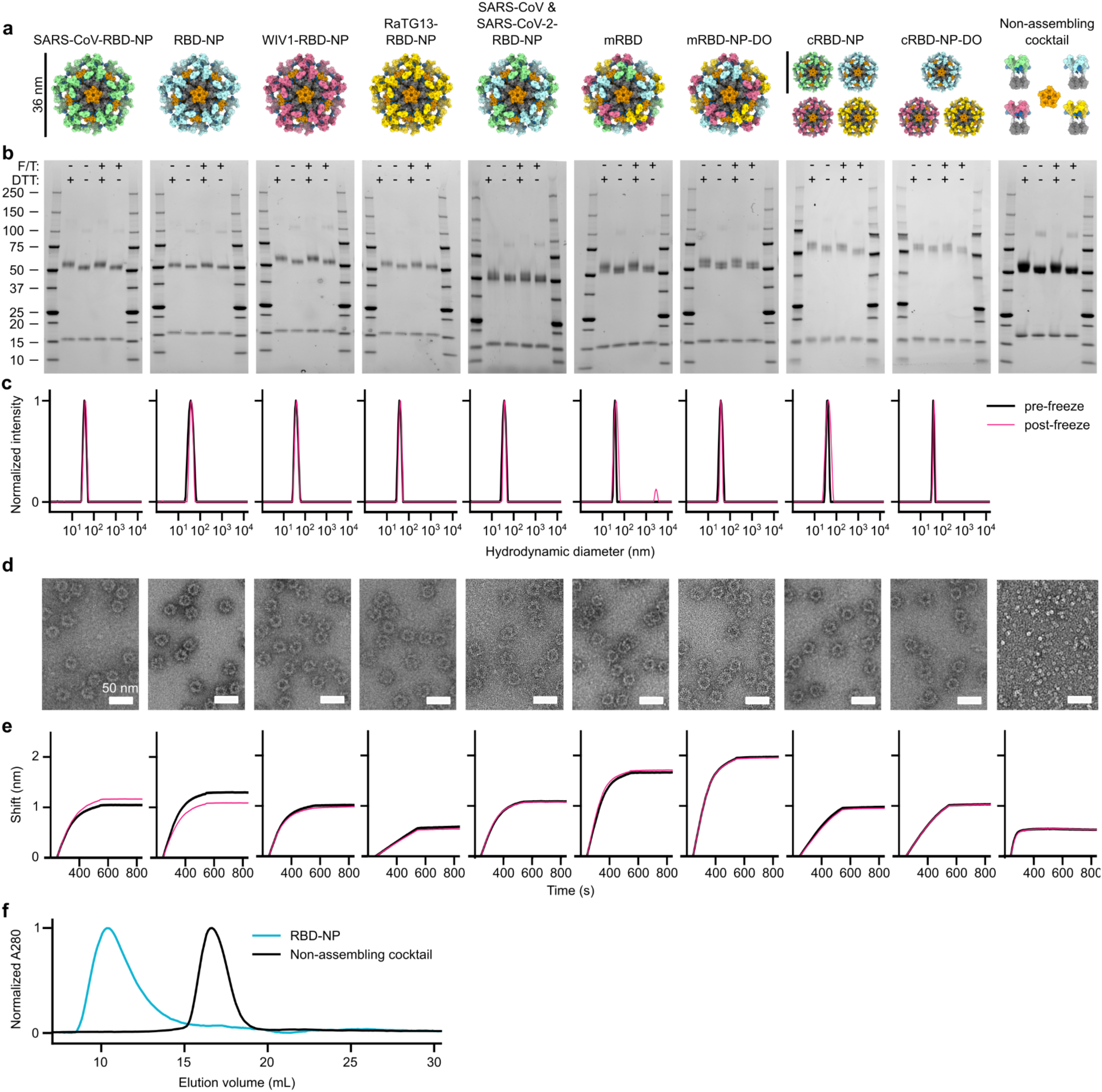
*In vitro* characterization of sarbecovirus RBD-NP immunogens. **a**, Designed models of the various vaccine candidates evaluated. Scale bars, 36 nm. **b**, SDS-PAGE analysis of purified nanoparticles. DTT, dithiothreitol; F/T, post-freeze/thaw. **c**, Dynamic light scattering. **d**, Electron micrographs of negatively stained samples. Scale bars, 50 nm. **e**, Binding of 100 nM SEC-purified nanoparticle immunogens and the non-assembling cocktail immunogen (which was not purified with SEC) to immobilized hACE2-Fc. **f**, SEC chromatogram overlay of purified RBD-NP and non-assembling cocktail.

**Extended Data Fig. 5:**
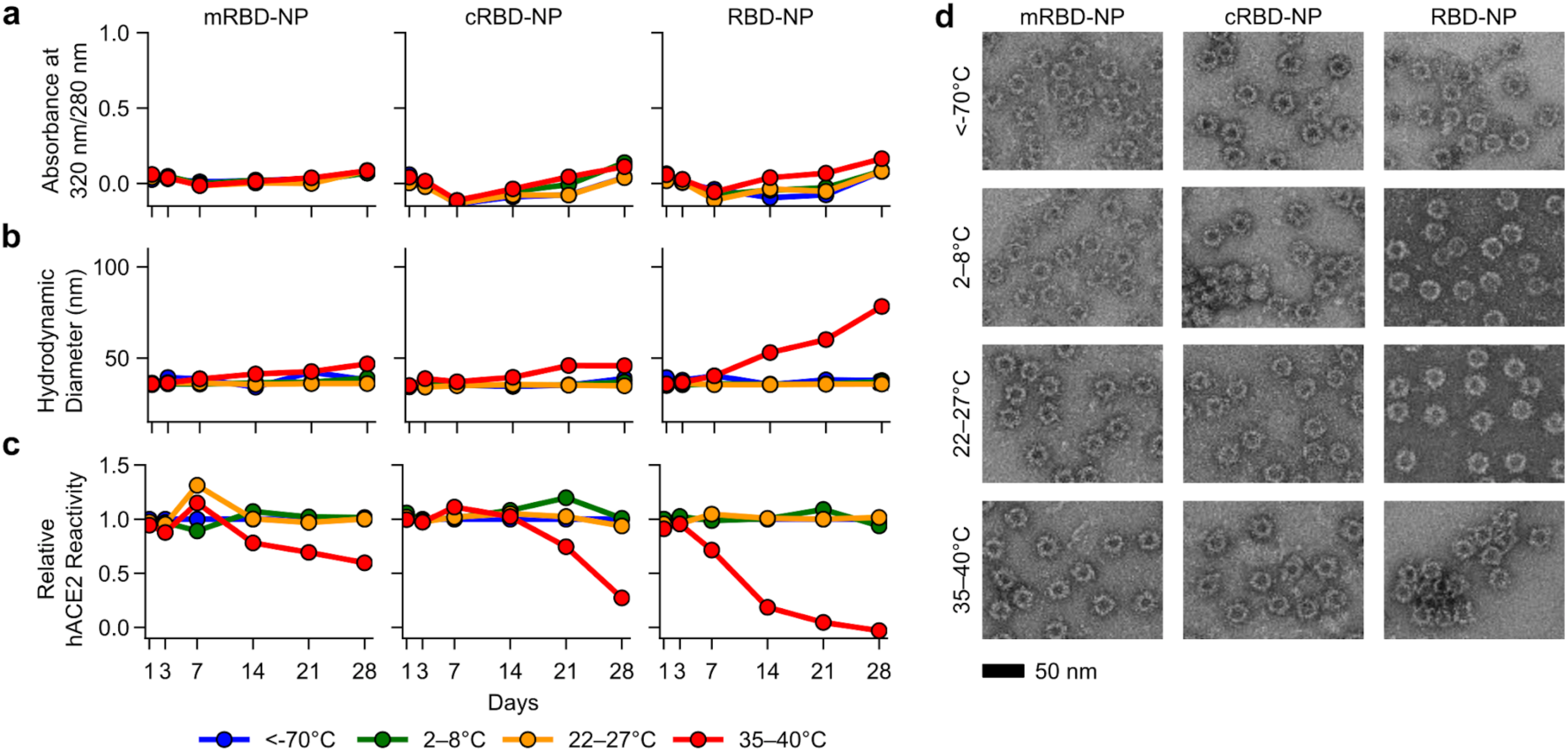
Accelerated stability studies of nanoparticle immunogens. The physical and antigenic stability of mRBD-NP, cRBD-NP, and (SARS-CoV-2) RBD-NP samples incubated at four different temperatures was followed for four weeks. **a**, The ratio of UV/vis absorbance at 320 nm/280 nm is a measure of turbidity. **b**, Hydrodynamic diameter of the nanoparticles measured using dynamic light scattering. **c**, hACE2 binding, measured by comparing the peak amplitude of hACE2 binding for each sample to a reference sample stored at <-70°C using biolayer interferometry. **d**, Electron micrographs of negatively stained samples after incubation for 28 days at the indicated temperatures. Scale bar, 50 nm.

**Extended Data Fig. 6:**
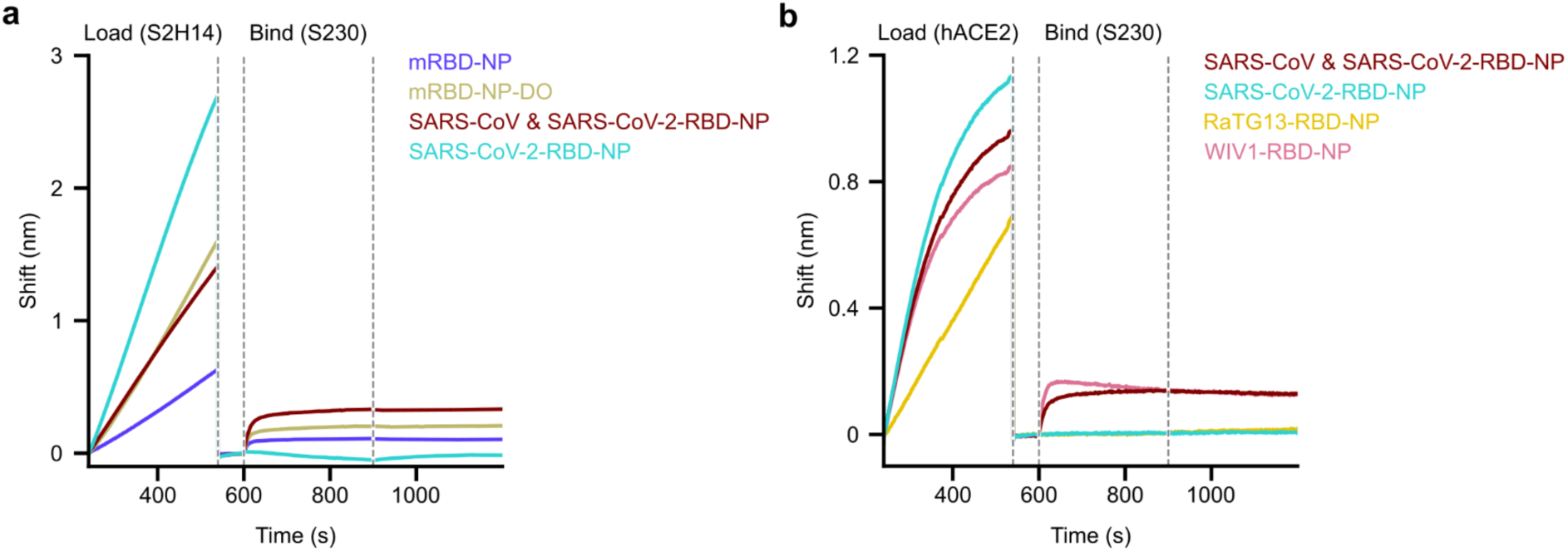
Confirmation of co-display using sandwich biolayer interferometry. **a**, The SARS-CoV-2 S-specific mAb S2H14 immobilized on protein A biosensors was used to capture various nanoparticle immunogens. The captured nanoparticles were subsequently exposed to a Fab derived from the SARS-CoV S-specific mAb S230. **b**, hACE2-Fc immobilized on protein A biosensors was used to capture various nanoparticle immunogens. The captured nanoparticles were subsequently exposed to a Fab derived from the SARS-CoV S-specific mAb S230.

### Mosaic RBD-NPs elicit cross-reactive and broadly neutralizing sarbecovirus Abs

The mosaic and cocktail nanoparticle immunogens were compared to monovalent RBD-NP vaccines and a non-assembling control vaccine comprising all four RBD-I53-50A trimeric components and a non-assembling I53-50B pentamer (**Extended Data Fig. 4**) in an immunization study in BALB/c mice (**Fig. 4a** and **Extended Data Fig. 7**). All immunizations comprised 1 μg of total RBD antigen, such that 0.25 μg of each RBD was given in each administration of the quadrivalent vaccines. After two immunizations, all four mosaic or cocktail RBD-NP vaccines elicited strong binding (GMT ∼5×10^4^) (**Extended Data Fig. 7a–d, f**) and potent serum neutralizing Ab titers against wild-type (D614G) SARS-CoV-2 S pseudovirus (GMT 2–3×10^3^) (**Fig. 4b**). The neutralizing Ab responses were ∼3-4-fold lower than that of the monovalent SARS-CoV-2 RBD-NP (8×10^3^), showing similar neutralization potency to the aforementioned low dose SARS-CoV-2 RBD-NP immunization study (**Fig. 2b)**. The neutralizing activity elicited by the other monovalent RBD-NPs and the non-assembling control were ∼2 orders of magnitude lower (GMT 0.4–2×10^2^) than for the monovalent SARS-CoV-2 RBD-NP. Although ELISA binding titers were comparable across mosaic and cocktail groups against SARS-CoV S (Urbani strain) (GMT 0.8–1×10^5^), the corresponding pseudovirus neutralization titers showed more nuanced patterns (**Fig. 4c** and **Extended Data Fig. 7 e,g**). Tetravalent mosaic and cocktail RBD-NPs elicited potent neutralizing activity (GMT 4×10^3^) with magnitudes roughly 3-fold lower than that of the monovalent SARS-CoV RBD-NP (GMT 1×10^4^). Strikingly, the trivalent nanoparticle immunogens (mRBD-NP-DO)—which did not contain the SARS-CoV RBD— also elicited potent neutralization (GMT ∼1×10^3^). This cross-neutralization likely arose from the inclusion of the WIV1 RBD (**Extended Data Fig. 7 h**) in the trivalent immunogens, as WIV1 cross-reacts with a SARS-CoV-specific mAb ^3^ (**Extended Data Fig. 7**) and the monovalent WIV1 RBD-NP induced similar levels of pseudovirus neutralization (GMT 2×10^3^) (**Fig. 4c**). The non-assembling control immunogen, which contains all four RBD-I53-50A trimeric components, also elicited substantial neutralizing activity against SARS-CoV (GMT ∼8×10^2^), but not against SARS-CoV-2. Finally, cRBD-NP, mRBD-NP-DO and especially mRBD-NP elicited higher serum neutralizing titers than the monovalent SARS-CoV-2 RBD-NP against the SARS-CoV-2 B.1.351 S HIV pseudotyped virus, indicating that this approach might enhance the breadth of Abs elicited and could overcome the emergence of variants of concern (**Fig. 4d**). These data show that both cRBD-NPs and mRBD-NPs are promising vaccine candidates for eliciting broad sarbecovirus immunity, in agreement with previous findings using a different nanoparticle platform ^76^ and recent results obtained with nanoparticle vaccines for influenza ^73^.

**Extended Data Fig 7:**
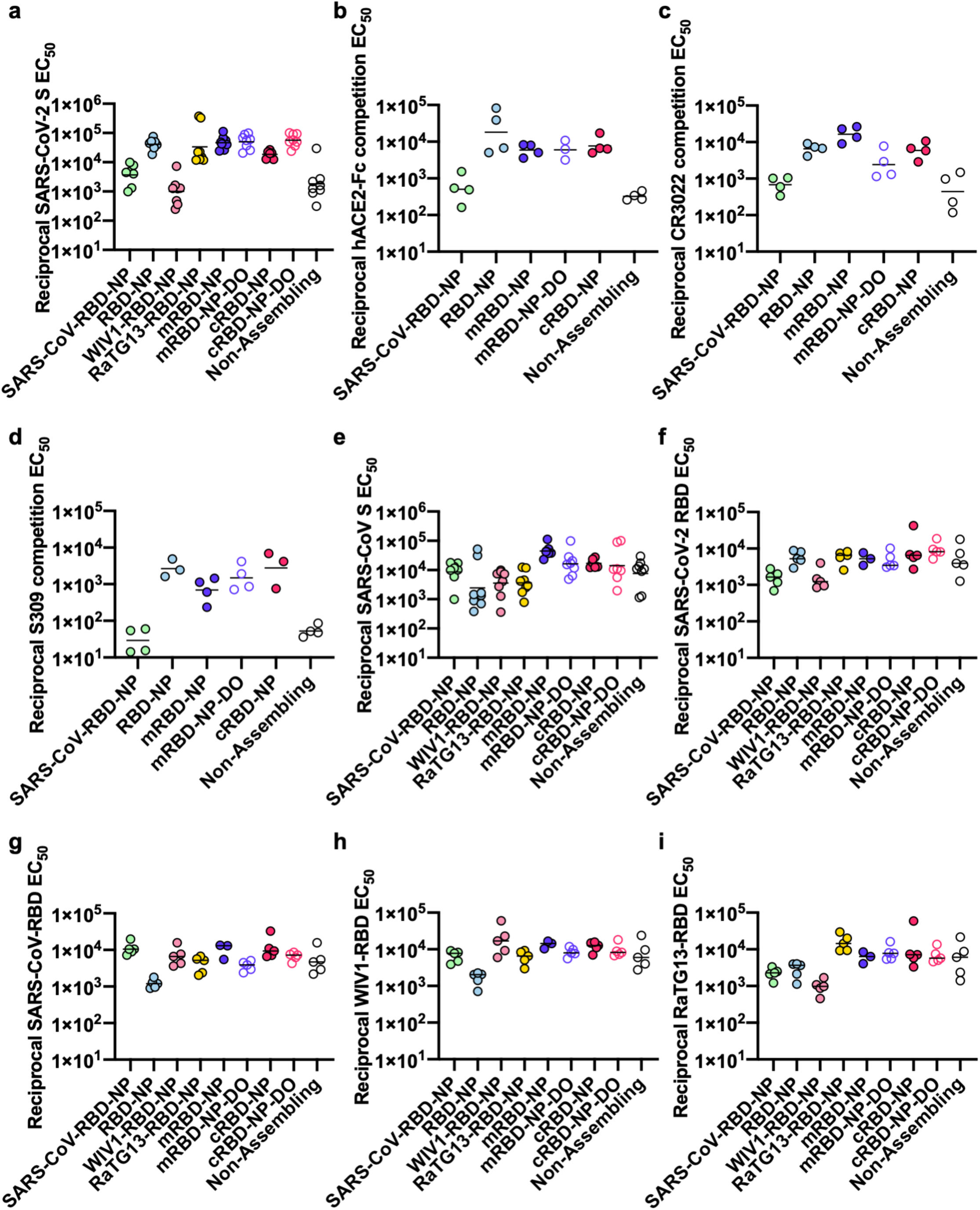
Serum Ab binding titers elicited by mosaic and cocktail RBD-NPs. (a) ELISA binding to SARS-CoV-2 S at week 5. (b–d) Titers of SARS-CoV-2 S-specific (b) ACE2-Fc (c) CR3022 (d) and S309 competing antibodies in immunized mouse sera. (e) ELISA binding to SARS-CoV S at week 5. (f–i) ELISA binding to RBDs from the (f) SARS-CoV-2 (g) SARS-CoV (h) WIV1 (i) RaTG13 spikes.

### Mosaic RBD-NPs protect mice against heterotypic SARS-CoV challenge

To gauge the ability of the multivalent RBD-NPs to confer protection against vaccine-matched and heterotypic sarbecoviruses, we challenged 6 mice from several of the groups with a high dose (10^5^ pfu) of the mouse-adapted SARS-CoV MA15 virus ^80^. In agreement with the pseudovirus neutralization data, animals immunized with the monovalent SARS-CoV RBD-NP, tetravalent mRBD-NP, and the non-assembling control immunogen were protected from weight loss and serious lung pathology throughout the 4 days of the experiment. The animals receiving the monovalent SARS-CoV-2 RBD-NP experienced up to 12% average weight loss, whereas the unvaccinated mice exhibited further weight loss (up to 15%) and signs of lung pathology **(Fig. 4e–f)**. All but one mice vaccinated with the mosaic or cocktail nanoparticles were completely protected from viral replication in the lungs whereas we detected ∼10^3^ and 10^7^ pfu/lobe for half of the mice receiving the non-assembling control immunogen and all unvaccinated mice, respectively (**Fig. 4g**). Strikingly, the trivalent mRBD-NP-DO provided protection that was virtually indistinguishable from the tetravalent mRBD-NP, despite lacking the SARS-CoV RBD. These results provide proof-of-principle that mosaic and cocktail RBD nanoparticle vaccines elicit broad protection against heterotypic sarbecovirus challenge and could represent a next generation of vaccines developed in anticipation of future spillovers.

## Discussion

The data presented here show that a single immunization with SARS-CoV-2 RBD-NP confers protection against lethal SARS-CoV-2 MA10 challenge suggesting that the potent Ab responses elicited could enable dose-sparing regimens to achieve global vaccination. Moreover, SARS-CoV-2 RBD-NP, which is currently under evaluation in the clinic, elicits diverse Ab responses neutralizing a broad spectrum of SARS-CoV-2 variants detected in clinical isolates. The E484K mutation, however, leads to a ∼5-fold reduction in neutralizing activity elicited by either RBD-NP or HexaPro in NHPs. Although both SARS-CoV-2 RBD-NP and HexaPro-elicited sera robustly neutralize the B.1.1.7 S variant, which does not include the E484K substitution, neutralization of the B.1.351 variant was dampened, as was also the case with sera from individuals vaccinated twice with the Pfizer-BioNTech BNT162b2 mRNA. These findings are in agreement with a recent report showing that the serum neutralizing activity against the B.1.351 variant from mRNA-1273-vaccinated individuals was comparably reduced ^81^, as was also the case for neutralization of authentic SARS-CoV-2 B.1.351 by HCP ^43^. Collectively, these data indicate that a significant fraction of vaccine-elicited neutralizing activity is directed to the RBM, which is the target of potent neutralizing Abs, irrespective of the antigen design strategy (RBD- or S-based), the vaccine modality (protein subunit or mRNA), or species (NHPs or human).

The ongoing global spread of SARS-CoV-2 and the circulation of a large number of sarbecoviruses in bats ^3,4^ strongly motivate the development of vaccines that protect against a broad spectrum of coronaviruses. We observed that vaccination of NHPs with SARS-CoV-2 RBD-NP or HexaPro induced comparable but moderate neutralization breadth against genetically distinct sarbecoviruses. Both co-display (mosaic RBD-NPs) and co-immunization (cocktails of RBD-NPs) elicited robust neutralizing Ab responses against SARS-CoV-2 (including the B.1.351 variant of concern), SARS-CoV, and related pseudoviruses. Mice vaccinated with these multivalent vaccines were completely protected from disease upon severe SARS-CoV MA15 challenge, including with formulations that did not include the SARS-CoV RBD, highlighting the potential of this approach to achieve broad sarbecovirus immunity overcoming both the emergence of SARS-CoV-2 variants and putative future zoonosis of genetically distinct sarbecoviruses. Our results support the RBD-centric nature of neutralizing Ab responses resulting from infection and vaccination ^13,64,82^, irrespective of immunogen format or vaccine modality, and pave the way for advancing pan-sarbecovirus vaccines to the clinic.

**Data Table 1:**
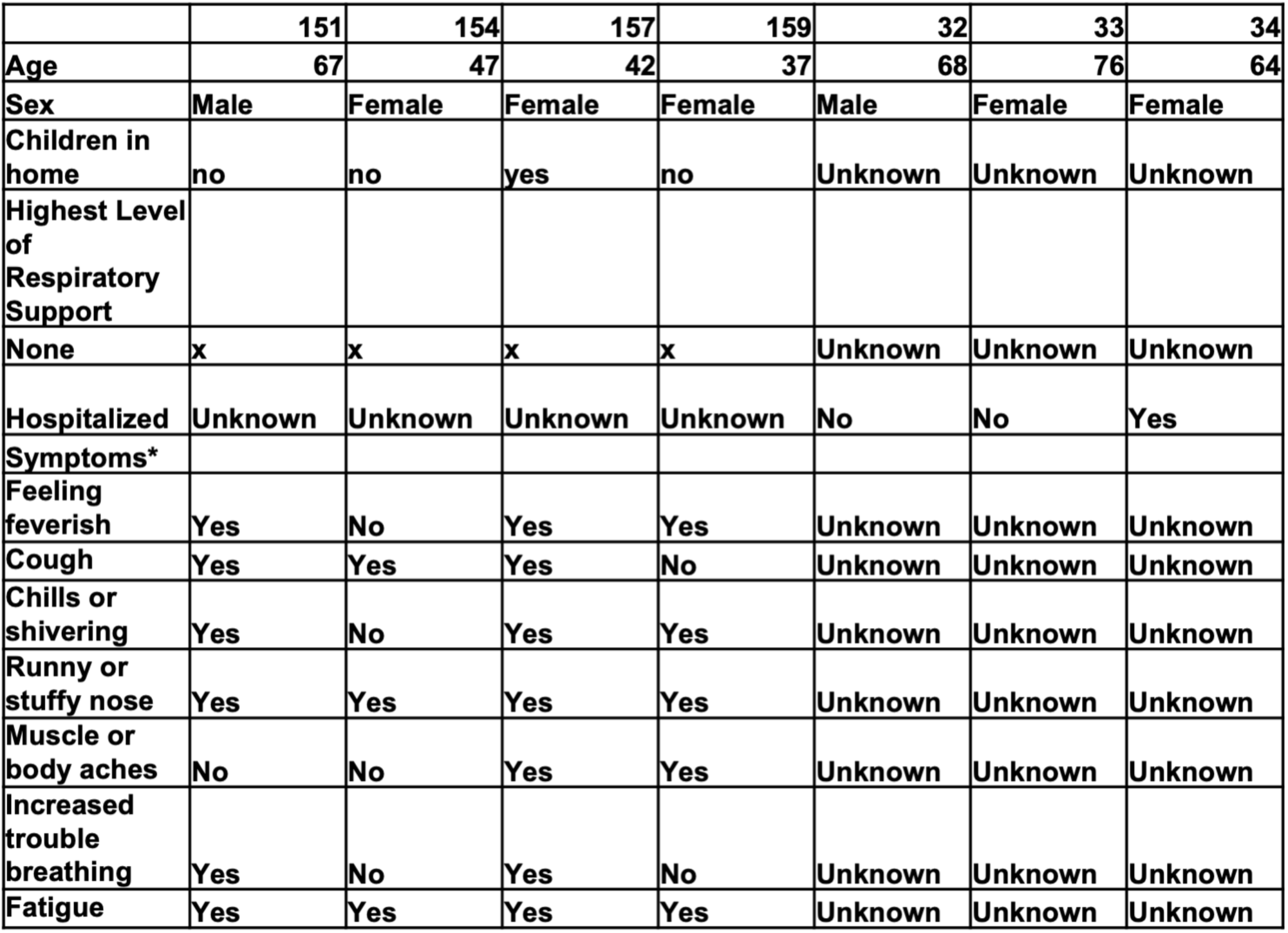
Human convalescent plasma samples.

## Acknowledgements

We thank Helen Chu and Marion Pepper for coordination of HCP samples and Ratika Krishnamurty for program management. This study was supported by the Bill & Melinda Gates Foundation (INV–017592 to J.S.M., OPP1156262 to N.P.K. and D.V.), a generous gift from the Audacious Project, a generous gift from Jodi Green and Mike Halperin, a generous gift from the Hanauer family, the Defense Threat Reduction Agency (HDTRA1-18-1-0001 to N.P.K.), the National Institute of General Medical Sciences (R01GM120553 to D.V.), the National Institute of Allergy and Infectious Diseases (R01-AI127521 to J.S.M, DP1AI158186 and HHSN272201700059C to D.V.), a Pew Biomedical Scholars Award (D.V.), Investigators in the Pathogenesis of Infectious Disease Awards from the Burroughs Wellcome Fund (D.V.), Fast Grants (D.V.) an Animal Models Contract HHSN272201700036I-75N93020F00001 (R.S.B), and the North Carolina Policy Collaboratory at the University of North Carolina at Chapel Hill with funding from the North Carolina Coronavirus Relief Fund established and appropriated by the North Carolina General Assembly (R.S.B).

## Author contributions

Conceptualization: A.C.W., P.S.A., H.K., B.P., N.P.K., D.V.; Modeling and design: A.C.W., N.P.K., D.V.; Formal Analysis: A.C.W., M.C.M., A.S., A.G., M.A.T., J.B., B.F., M.J.N., S.W., T.S., R.R., C.S., L.C., N.P.K, D.V.; Resources: M.A.O., P.S.A., S.W.T., R.R., D.T.O., R.V.D.M., W.V.W., D.C., C.-L.H., J.S.M., D.H.F., F.V., B.P.; Writing – Original Draft: A.C.W., N.P.K, D.V..; Writing – Review & Editing: All authors; Visualization: A.C.W., M.C.M., A.S., A.G., N.B., N.P.K., D.V.; Supervision: D.H.F., J.S.M., L.C., R.S.B., T.P.S., B.P., N.P.K., D.V.; Funding Acquisition: J.S.M., R.S.B., N.P.K., D.V.

## Declaration of interest

A.C.W, N.P.K. and D.V., are named as inventors on patent applications filed by the University of Washington based on the studies presented in this paper. N.P.K. is a co-founder, shareholder, paid consultant, and chair of the scientific advisory board of Icosavax, Inc. and has received an unrelated sponsored research agreement from Pfizer. D.V. is a consultant for and has received an unrelated sponsored research agreement from Vir Biotechnology Inc. R.R., D.T.O., and R.V.D.M are employees of GlaxoSmithKline. C.-L.H. and J.S.M. are inventors on U.S. patent application no. 63/032,502 “Engineered Coronavirus Spike (S) Protein and Methods of Use Thereof. The other authors declare no competing interests.

## Methods

### Cell lines

Expi293F cells are derived from the HEK293F cell line (Life Technologies). Expi293F cells were grown in Expi293 Expression Medium (Life Technologies), cultured at 36.5°C with 8% CO_2_ and shaking at 150 rpm. HEK293T/17 is a female human embryonic kidney cell line (ATCC). The HEK-ACE2 adherent cell line was obtained through BEI Resources, NIAID, NIH: Human Embryonic Kidney Cells (HEK293T) Expressing Human Angiotensin-Converting Enzyme 2, HEK293T-hACE2 Cell Line, NR-52511. All adherent cells were cultured at 37°C with 8% CO_2_ in flasks with DMEM + 10% FBS (Hyclone) + 1% penicillin-streptomycin. Cell lines other than Expi293F were not tested for mycoplasma contamination nor authenticated.

### Mice

Female BALB/c mice (Stock # 000651, Balb/c cByJ mice) four weeks old were obtained from Jackson Laboratory, Bar Harbor, Maine, and maintained at the Comparative Medicine Facility at the University of Washington, Seattle, WA, accredited by the American Association for the Accreditation of Laboratory Animal Care International (AAALAC). Animal procedures were performed under the approvals of the Institutional Animal Care and Use Committee (IACUC) of University of Washington, Seattle, WA, and University of North Carolina, Chapel Hill, NC.

### Pigtail macaques

Two adult male Pigtail macaques (*Macaca nemestrina*) were immunized in this study. All animals were housed at the Washington National Primate Research Center (WaNPRC), an AAALAC International accredited institution. All experiments were approved by The University of Washington’s IACUC. Animals were singly housed in comfortable, clean, adequately-sized cages with ambient temperatures between 72–82°F. Animals received environmental enrichment for the duration of the study including grooming contact, perches, toys, foraging experiences and access to additional environment enrichment devices. Water was available through automatic watering devices and animals were fed a commercial monkey chow, supplemented daily with fruits and vegetables. Throughout the study, animals were checked twice daily by husbandry staff.

### Rhesus macaques

Adapted from Arunachalam et al. 2021. Rhesus macaques (Macaca mulatta) of Indian origin, aged 3–7 years were assigned to the study (Arunachalam et al. 2021). Animals were distributed between the groups such that the age and weight distribution were comparable across the groups. Animals were housed and maintained at the New Iberia Research Center (NIRC) of the University of Louisiana at Lafayette, an AAALAC International accredited institution, in accordance with the rules and regulations of the Guide for the Care and Use of Laboratory Animal Resources. The entire study (protocol 2020-8808-15) was reviewed and approved by the University of Louisiana at Lafayette IACUC. All animals were negative for SIV, simian T cell leukemia virus, and simian retrovirus.

### Convalescent human sera

Samples collected between 1–60 days post infection from individuals who tested positive for SARS-CoV-2 by PCR were profiled for anti-SARS-CoV-2 S antibody responses and those with anti-S Ab responses were maintained in the cohort ^83^. Individuals were enrolled as part of the HAARVI study at the University of Washington in Seattle, WA. Baseline sociodemographic and clinical data for these individuals are summarized in Data 1. This study was approved by the University of Washington Human Subjects Division Institutional Review Board (STUDY00000959 and STUDY00003376). All experiments were performed in at least two replicates. One sample is the 20/130 COVID-19 plasma from NIBSC (https://www.nibsc.org/documents/ifu/20-130.pdf).

### Pfizer vaccinated human sera

Blood samples were collected from participants who had received both doses of the Pfizer mRNA vaccine and were 7–30 days post second vaccine dose. Individuals were enrolled in the UWARN: COVID-19 in WA study at the University of Washington in Seattle, WA. This study was approved by the University of Washington Human Subjects Division Institutional Review Board (STUDY00010350).

## Method Details

### Plasmid construction

The SARS-CoV-2-RBD-Avi construct was synthesized by GenScript into pcDNA3.1- with an N-terminal mu-phosphatase signal peptide and a C-terminal octa-histidine tag, flexible linker, and avi tag (GHHHHHHHHGGSSGLNDIFEAQKIEWHE). The boundaries of the construct are N-_328_RFPN_331_ and _528_KKST_531_-C (Walls et al., 2020). The SARS-CoV-2-B.1.351-RBD-Avi was synthesized by GenScript into CMVR with K417N, E484K, and N501Y mutations from the Wuhan-1 strain used in the SARS-CoV-2-RBD-Avi construct listed above. The GD-Pangolin (326-527), WIV1 (316-518), RaTG13 (359-562), and ZXC21 (323-507) were synthesized by GenScript into vector pcDNA3.1- with a preceding mu-phosphatase signal peptide and a C-terminal octahistidine tag, flexible linker, and avi tag (GHHHHHHHHGGSSGLNDIFEAQKIEWHE). SARS-CoV-1 (306-575) was subcloned from a GenArt synthesized SARS-CoV-1 Spike ectodomain. The SARS-CoV S2P was synthesized by GeneArt and placed into a modified pOPING vector with its original <u>N-terminal</u> mu-phosphatase <u>signal peptide</u>, and an engineered <u>C-terminal</u> extension: SG-RENLYFQG (TEV *<u>protease</u>* site), GGGSG-YIPEAPRDGQAYVRKDGEWVLLSTFL (foldon trimerization motif), G-HHHHHH (hexa-histidine tag), just upstream of the predicted transmembrane region (YIK). The SARS-CoV S was stabilized with the 2P mutations (Pallesen et al., 2017). The SARS-CoV-2 S-2P ectodomain trimer (GenBank: <u>YP_009724390.1</u>, BEI NR-52420) was synthesized by GenScript into pCMV with an N-terminal mu-phosphatase signal peptide and a C-terminal TEV cleavage site (GSGRENLYPQG), T4 fibritin foldon (GGGSGYIPEAPRDGQAYVRKDGEWVLLSTPL), and octa-histidine tag (GHHHHHHHH) (Walls et al., 2020). The construct contains the 2P mutations (proline substitutions at residues 986 and 987; (Pallesen et al., 2017)) and an _682_SGAG_685_ substitution at the furin cleavage site. The SARS-CoV-2 RBD was genetically fused to the N terminus of the trimeric I53-50A nanoparticle component using 12 or 16 glycine and serine residues. RBD-12GS-I53-50A fusions were synthesized and cloned by Genscript into pCMV. The RBD-16GS-I53-50A fusion was cloned into pCMV/R using the Xba1 and AvrII restriction sites and Gibson assembly (<u>Gibson et al., 2009</u>). All RBD-bearing components contained an N-terminal mu-phosphatase signal peptide and a C-terminal octa-histidine tag. Human ACE2 ectodomain was genetically fused to a sequence encoding a thrombin cleavage site and a human Fc fragment at the C-terminal end. hACE2-Fc was synthesized and cloned by GenScript with a BM40 signal peptide. CR3022 Genes encoding CR3022 heavy and light chains were ordered from GenScript and cloned into pCMV/R. Antibodies were expressed by transient co-transfection of both heavy and light chain plasmids. S309 construct as previously described (Pinto et al. 2020). SARS-CoV-2 Hexapro construct is as previously described (Hsieh et al.) and placed into CMVR with an octa-his tag. Plasmids were transformed into the NEB 5-alpha strain of *E. coli* (New England Biolabs) for subsequent DNA extraction from bacterial culture (NucleoBond Xtra Midi kit) to obtain plasmid for transient transfection into Expi293F cells. The amino acid sequences of all novel proteins used in this study can be found in <u>Data S1</u>.

### Transient transfection

Proteins were produced in Expi293F cells grown in suspension using Expi293F expression medium (Life Technologies) at 33°C, 70% humidity, 8% CO_2_ rotating at 150 rpm. The cultures were transfected using PEI-MAX (Polyscience) with cells grown to a density of 3.0 million cells per mL and cultivated for 3 days. Supernatants were clarified by centrifugation (5 min at 4000 rcf), addition of PDADMAC solution to a final concentration of 0.0375% (Sigma Aldrich, #409014), and a second spin (5 min at 4000 rcf).

### Microbial protein expression and purification

The I53-50A and I53-50B.4.PT1 proteins ^48^ were expressed in Lemo21(DE3) (NEB) in LB (10 g Tryptone, 5 g Yeast Extract, 10 g NaCl) grown in 2 L baffled shake flasks or a 10 L BioFlo 320 Fermenter (Eppendorf). Cells were grown at 37°C to an OD600 ∼0.8, and then induced with 1 mM IPTG. Expression temperature was reduced to 18°C and the cells shaken for ∼16 h. The cells were harvested and lysed by microfluidization using a Microfluidics M110P at 18,000 psi in 50 mM Tris, 500 mM NaCl, 30 mM imidazole, 1 mM PMSF, 0.75% CHAPS. Lysates were clarified by centrifugation at 24,000 g for 30 min and applied to a 2.6×10 cm Ni Sepharose 6 FF column (Cytiva) for purification by IMAC on an AKTA Avant150 FPLC system (Cytiva). Protein of interest was eluted over a linear gradient of 30 mM to 500 mM imidazole in a background of 50 mM Tris pH 8, 500 mM NaCl, 0.75% CHAPS buffer. Peak fractions were pooled, concentrated in 10K MWCO centrifugal filters (Millipore), sterile filtered (0.22 μm) and applied to either a Superdex 200 Increase 10/300, or HiLoad S200 pg GL SEC column (Cytiva) using 50 mM Tris pH 8, 500 mM NaCl, 0.75% CHAPS buffer. I53-50A elutes at ∼0.6 column volume (CV). I53-50B.4PT1 elutes at ∼0.45 CV. After sizing, bacterial-derived components were tested to confirm low levels of endotoxin before using for nanoparticle assembly.

### Protein purification

Proteins containing His tags were purified from clarified supernatants via a batch bind method where each clarified supernatant was supplemented with 1 M Tris-HCl pH 8.0 to a final concentration of 45 mM and 5 M NaCl to a final concentration of ∼310 mM. Talon cobalt affinity resin (Takara) was added to the treated supernatants and allowed to incubate for 15 min with gentle shaking. Resin was collected using vacuum filtration with a 0.2 μm filter and transferred to a gravity column. The resin was washed with 20 mM Tris pH 8.0, 300 mM NaCl, and the protein was eluted with 3 column volumes of 20 mM Tris pH 8.0, 300 mM NaCl, 300 mM imidazole. The batch bind process was then repeated and the first and second elutions combined. SDS-PAGE was used to assess purity. RBD-I53-50A fusion protein IMAC elutions were concentrated to > 1 mg/mL and subjected to three rounds of dialysis into 50 mM Tris pH 7.4, 185 mM NaCl, 100 mM Arginine, 4.5% glycerol, and 0.75% w/v 3-[(3-cholamidopropyl)dimethylammonio]-1-propanesulfonate (CHAPS) in a hydrated 10K molecular weight cutoff dialysis cassette (Thermo Scientific). S-2P and Hexapro IMAC elution fractions were concentrated to ∼1 mg/mL and dialyzed three times into 50 mM Tris pH 8, 150 mM NaCl, 0.25% L-Histidine in a hydrated 10K molecular weight cutoff dialysis cassette (Thermo Scientific). Due to inherent instability, the S-2P trimer was immediately flash frozen and stored at −80°C.

Clarified supernatants of cells expressing monoclonal antibodies and human ACE2-Fc were purified using a MabSelect PrismA 2.6 × 5 cm column (Cytiva) on an AKTA Avant150 FPLC (Cytiva). Bound antibodies were washed with five column volumes of 20 mM NaPO_4_, 150 mM NaCl pH 7.2, then five column volumes of 20 mM NaPO_4_, 1 M NaCl pH 7.4 and eluted with three column volumes of 100 mM glycine at pH 3.0. The eluate was neutralized with 2 M Trizma base to 50 mM final concentration. SDS-PAGE was used to assess purity.

Recombinant S309 was expressed as a Fab in expiCHO cells transiently co-transfected with plasmids expressing the heavy and light chain, as described above (see Transient transfection) (<u>Stettler et al., 2016</u>). The protein was affinity-purified using a HiTrap Protein A Mab select Xtra column (Cytiva) followed by desalting against 20 mM NaPO_4_, 150 mM NaCl pH 7.2 using a HiTrap Fast desalting column (Cytiva). The protein was sterilized with a 0.22 μm filter and stored at 4°C until use.

### *In vitro* nanoparticle assembly and purification

Total protein concentration of purified individual nanoparticle components was determined by measuring absorbance at 280 nm using a UV/vis spectrophotometer (Agilent Cary 8454) and calculated extinction coefficients (<u>Gasteiger et al., 2005</u>). The assembly steps were performed at room temperature with addition in the following order: RBD-I53-50A trimeric fusion protein, followed by additional buffer (50 mM Tris pH 7.4, 185 mM NaCl, 100 mM Arginine, 4.5% glycerol, and 0.75% w/v CHAPS) as needed to achieve desired final concentration, and finally I53-50B.4PT1 pentameric component (in 50 mM Tris pH 8, 500 mM NaCl, 0.75% w/v CHAPS), with a molar ratio of RBD-I53-50A:I53-50B.4PT1 of 1.1:1. All RBD-I53-50 *in vitro* assemblies were incubated at 2-8°C with gentle rocking for at least 30 min before subsequent purification by SEC in order to remove residual unassembled component. Different columns were utilized depending on purpose: Superose 6 Increase 10/300 GL column was used analytically for nanoparticle size estimation, a Superdex 200 Increase 10/300 GL column used for small-scale pilot assemblies, and a HiLoad 26/600 Superdex 200 pg column used for nanoparticle production. Assembled particles were purified in 50 mM Tris pH 7.4, 185 mM NaCl, 100 mM Arginine, 4.5% glycerol, and 0.75% w/v CHAPS, and elute at ∼11 mL on the Superose 6 column and in the void volume of Superdex 200 columns. Assembled nanoparticles were sterile filtered (0.22 μm) immediately prior to column application and following pooling of fractions.

### Endotoxin measurements

Endotoxin levels in protein samples were measured using the EndoSafe Nexgen-MCS System (Charles River). Samples were diluted 1:50 or 1:100 in Endotoxin-free LAL reagent water, and applied into wells of an EndoSafe LAL reagent cartridge. Charles River EndoScan-V software was used to analyze endotoxin content, automatically back-calculating for the dilution factor. Endotoxin values were reported as EU/mL which were then converted to EU/mg based on UV/vis measurements. Our threshold for samples suitable for immunization was < 50 EU/mg.

### Pigtail macaque immunization

Two Pigtail macaques were immunized with 250 μg of RBD-12GS-I53-50 nanoparticle (88 μg RBD antigen) with AddaVax at day 0, day 28, and O/W at day 168. Blood was collected every 14 days post-prime. Blood was collected in serum collection tubes and allowed to clot at room temperature. Serum was isolated after a 15 min spin at 1455 x g for 15 min and stored at −80C until use. Prior to vaccination or blood collection, animals were sedated with an intramuscular injection (10 mg/kg) of ketamine (Ketaset®; Henry Schein). Prior to inoculation, immunogen suspensions were gently mixed 1:1 vol/vol with AddaVax adjuvant for immunizations 1 and 2 and O/W for immunization 3 (Invivogen, San Diego, CA) to reach a final concentration of 0.250 mg/mL antigen. The vaccine was delivered intramuscularly into both quadriceps muscles with 1 mL per injection site on days 0, 28, and 168. All injection sites were shaved prior to injection. Animals were observed daily for general health (activity and appetite, urine/feces output) and for evidence of reactogenicity at the vaccine inoculation site (swelling, erythema, and pruritus) for up to 1 week following vaccination. They also received physical exams including temperature and weight measurements at each study time point. None of the animals became severely ill during the course of the study nor required euthanasia.

### Rhesus macaque immunization

Adapted from ^50^. AS03 was kindly provided by GSK Vaccines. AS03 is an oil-in-water emulsion that contains 11.86 mg a-tocopherol, 10.69 mg squalene, and 4.86 mg polysorbate 80 (Tween-80) in PBS. For each dose, RBD-NP was diluted to 50 μg/ml (RBD component) in 250 μl of Tris-buffered saline (TBS) and mixed with an equal volume of AS03.The dose of AS03 was 50% v/v (equivalent of one human dose). Soluble Hexapro was diluted to 50 μg/ml in 250 μl of Tris-buffered saline (TBS) and mixed with an equal volume of AS03. All immunizations were administered via the intramuscular route in right forelimbs. The volume of each dose was 0.5 ml.

### Deep Mutational Scanning

All mutations that escape serum antibody binding were mapped via a deep mutational scanning approach ^38,64^. We used previously described yeast-display RBD mutant libraries ^38,62^. Briefly, duplicate mutant libraries were constructed in the spike receptor binding domain (RBD) from SARS-CoV-2 (isolate Wuhan-Hu-1, Genbank accession number MN908947, residues N331-T531) and contain 3,804 of the 3,819 possible amino-acid mutations, with >95% present as single mutants. Each RBD variant was linked to a unique 16-nucleotide barcode sequence to facilitate downstream sequencing. As previously described, libraries were sorted for RBD expression and ACE2 binding to eliminate RBD variants that are completely misfolded or non-functional (i.e., lacking modest ACE2 binding affinity) ^38^.

Antibody escape mapping experiments were performed in biological duplicate using two independent mutant RBD libraries, with minor modifications from ^38^, and exactly as described in ^64^. The antibody escape mapping for the vaccinated NHP serum was performed in this study; the antibody escape mapping from convalescent human plasma was performed in ^64^. Briefly, mutant yeast libraries induced to express RBD were washed and incubated with plasma or serum at a range of dilutions for 1 h at room temperature with gentle agitation. For each sample, we chose a sub-saturating dilution such that the amount of fluorescent signal due to plasma antibody binding to RBD was approximately equal across samples. A 1:1000 dilution was used for the vaccinated NHP serum, and the exact dilutions of human convalescent plasma are reported in ^64^. After the antibody incubations, the libraries were secondarily labeled with 1:100 FITC-conjugated anti-MYC antibody (Immunology Consultants Lab, CYMC-45F) to label for RBD expression and and 1:200 Alexa-647-conjugated goat anti-human-IgA+IgG+IgM (Jackson ImmunoResearch 109-605-064) to label for bound serum or plasma antibodies. A flow cytometric selection gate was drawn to capture approximately 5% of the RBD mutant libraries with the lowest amount of plasma binding for their degree of RBD expression. For each sample, approximately 10 million RBD+ cells were processed on the cytometer. Antibody-escaped cells were grown overnight in SD-CAA (6.7g/L Yeast Nitrogen Base, 5.0g/L Casamino acids, 1.065 g/L MES acid, and 2% w/v dextrose) to expand cells prior to plasmid extraction.

Plasmid samples were prepared from pre-selection and overnight cultures of antibody-escaped cells (Zymoprep Yeast Plasmid Miniprep II) as previously described ^38^. The 16-nucleotide barcode sequences identifying each RBD variant were amplified by PCR and sequenced on an Illumina HiSeq 2500 with 50 bp single-end reads as described in ^38,62^.

Escape fractions were computed as described in ^38^, and exactly as described in ^64^. We used the `dms_variants` package (https://jbloomlab.github.io/dms_variants/, version 0.8.2) to process Illumina sequences into counts of each barcoded RBD variant in each pre-sort and antibody-escape population using the barcode/RBD look-up table from ^84^.

For each serum selection, we computed the “escape fraction” for each barcoded variant using the deep sequencing counts for each variant in the original and serum-escape populations and the total fraction of the library that escaped antibody binding via the formula provided in ^38^. These escape fractions represent the estimated fraction of cells expressing that specific variant that fall in the antibody escape bin, such that a value of 0 means the variant is always bound by serum and a value of 1 means that it always escapes serum binding. We then applied a computational filter to remove variants with low sequencing counts or highly deleterious mutations that might cause antibody escape simply by leading to poor expression of properly folded RBD on the yeast cell surface ^38,62^. Specifically, we removed variants that had (or contained mutations with) ACE2 binding scores < −2.35 or expression scores < −1, using the variant-and mutation-level deep mutational scanning scores from ^62^. Note that these filtering criteria are slightly more stringent than those used in ^38^ but are identical to those used in ^14,84^.

We next deconvolved variant-level escape scores into escape fraction estimates for single mutations using global epistasis models ^85^ implemented in the `dms_variants` package, as detailed at (https://jbloomlab.github.io/dms_variants/dms_variants.globalepistasis.html) and described in ^38^. The reported escape fractions throughout the paper are the average across the libraries (correlations shown in Extended Data 1); these scores are also in Supplementary Table 1 and at https://github.com/jbloomlab/SARS-CoV-2-RBD_MAP_RBD-nano-vax-NHP1/blob/main/results/supp_data/NHP_HCS_raw_data.csv. Sites of strong escape from each antibody were determined heuristically as sites whose summed mutational escape scores were at least 10 times the median sitewise sum of selection, and within 10-fold of the sitewise sum of the most strongly selected site. Sites shown in Figure 1a and Supplementary Figure 1a are the sites of strong escape for any of the three human convalescent plasma, plus sites 417, 452, and 501 due to their prevalence in circulating SARS-CoV-2 variants. For each plasma, the y-axis is scaled to be the greatest of (a) the maximum site-wise escape metric observed for that plasma, (b) 20x the median site-wise escape fraction observed across all sites for that plasma, or (c) an absolute value of 1.0 (to appropriately scale plasma that are not “noisy” but for which no mutation has a strong effect on plasma binding). Full documentation of the computational analysis is at https://github.com/jbloomlab/SARS-CoV-2-RBD_MAP_RBD-nano-vax-NHP1. These results are also available in a zoomable, interactive form at https://jbloomlab.github.io/SARS-CoV-2-RBD_MAP_RBD-nano-vax-NHP1/.

### ELISA

For anti-S-2P ELISA, 25 μL of 2 μg/mL S-2P was plated onto 384-well Nunc Maxisorp (ThermoFisher) plates in PBS and sealed overnight at RT. The next day plates were washed 4 × in Tris Buffered Saline Tween (TBST) using a plate washer (BioTek) and blocked with SuperBlock (ThermoFisher) for 1 h at 37°C. Plates were washed 4 × in TBST and 1:5 serial dilutions of mouse, NHP, or human sera were made in 25 μL TBST and incubated at 37°C for 1 h. Plates were washed 4 × in TBST, then anti-mouse (Invitrogen), anti-NHP (AlphaDiagnostics), or anti-human (Invitrogen) horseradish peroxidase-conjugated antibodies were diluted 1:5,000 and 25 μL added to each well and incubated at 37°C for 1 h. Plates were washed 4 × in TBST and 25 μL of TMB (SeraCare) was added to every well for 5 min at room temperature. The reaction was quenched with the addition of 25 μL of 1 N HCl. Plates were immediately read at 450 nm on a BioTek plate reader and data plotted and fit in Prism (GraphPad) using nonlinear regression sigmoidal, 4PL, X is log(concentration) to determine EC_50_ values from curve fits.

### Competition ELISA of NHP sera with hACE2, CR3022 IgG, and S309 IgGfor immobilized SARS-CoV-2 S2P

384-well Maxisorp plates (Thermo Fisher) were coated overnight at room temperature with 3 µg/mL of SARS-CoV-2 S2P (Pallesen et al. 2017) in 20mM Tris pH 8 and 150mM NaCl. Plates were slapped dry and blocked with Blocker Casein in TBS (Thermo Fisher) for one hour at 37°C. Plates were slapped dry and NHP sera was serially diluted 1:4 in TBST with an initial dilution of 1:4 for hACE2 competition or 1:2 for antibody competition. Random primary amine biotinylated (Thermo Fisher) hACE2-Fc, CR3022 (Yuan and Wu et al. 2020), or S309 (Pinto et al. 2020) were added, bringing the concentration of each well to the EC_50_ values of 0.2nM, 2nm, and 0.01nM, respectively. Plates were left for one hour at 37°C, then washed 4x with TBST using a 405 TS Microplate Washer (BioTek) followed by addition of 1:500 streptavidin-HRP (Thermo Fisher) for one hour at 37°C. Plates were washed 4x and TMB Microwell Peroxidase (Seracare) was added. The reaction was quenched after 1-2 minutes with 1 N HCl and the A450 of each well was read using a BioTek plate reader (BioTek).

### Competition ELISA of ---mouse sera and immobilized hACE2 or antibodies for SARS-CoV-2 S2P or SARS-CoV S2P

384-well Maxisorp plates (Thermo Fisher) were coated overnight at room temperature with 3 µg/mL of hACE2-Fc, CR3022 (Yuan and Wu et al. 2020), or S309 (Pinto et al. 2020) in 20mM Tris pH 8 and 150mM NaCl. Plates were slapped dry and blocked with Blocker Casein in TBS (Thermo Fisher) for one hour at 37°C. Plates were slapped dry and a 30-minute pre-incubated 1:5 serial dilution of mouse sera in TBST, with in initial dilution of 1:50 for hACE2-Fc competition or 1:10 for antibody competition, and a constant concentration of biotinylated (Avidity) SARS-CoV-2 S2P or SARS-CoV 2P at their EC_50_ values were added. Spike concentrations were 0.63nM, 5.98nM, and 0.22nM of SARS-CoV-2 S2P or 4.11nM, 2.89nM, and 0.19nM of SARS-CoV S2P for immobilized hACE2, CR3022, and S309, respectively. Plates were left for one hour at 37°C, then washed 4x with TBST using a 405 TS Microplate Washer (BioTek) followed by addition of 1:500 streptavidin-HRP (Thermo Fisher) for one hour at 37°C. Plates were washed 4x and TMB Microwell Peroxidase (Seracare) was added. The reaction was quenched after 1-2 minutes with 1 N HCl and the A450 of each well was read using a BioTek plate reader (BioTek).

### Pseudovirus Production

MLV-based D614G SARS-CoV-2 S, SARS-CoV-2-B-1.351, SARS-CoV S, and WIV-1 S pseudotypes were prepared as previously described (Millet and Whittaker, 2016; Walls et al., 2020 a+b). Briefly, HEK293T cells were co-transfected using Lipofectamine 2000 (Life Technologies) with an S-encoding plasmid, an MLV Gag-Pol packaging construct, and the MLV transfer vector encoding a luciferase reporter according to the manufacturer’s instructions. Cells were washed 3 × with Opti-MEM and incubated for 5 h at 37°C with transfection medium. DMEM containing 10% FBS was added for 60 h. The supernatants were harvested by spinning at 2,500 g, filtered through a 0.45 μm filter, concentrated with a 100 kDa membrane for 10 min at 2,500 g and then aliquoted and stored at −80°C. HIV-based pseudotypes were prepared as previously described (Crawford et al 2020). Briefly, HEK293T cells were cotransfected using Lipofectamine 2000 (Life Technologies) with an S-encoding plasmid, an HIV Gag-Pol, Tat, Rev1B packaging construct, and the HIV transfer vector encoding a luciferase reporter according to the manufacturer’s instructions. Cells were washed 3 × with Opti-MEM and incubated for 5 h at 37°C with transfection medium. DMEM containing 10% FBS was added for 60 h. The supernatants were harvested by spinning at 2,500 g, filtered through a 0.45 μm filter, concentrated with a 100 kDa membrane for 10 min at 2,500 g and then aliquoted and stored at −80°C.

D614G SARS-CoV-2 S (YP 009724390.1), RaTG13 S (QHR63300.2), Pangolin-Guangdong S (QLR06867.1), SARS-CoV S (YP 009825051.1), WIV1 S (AGZ48831.1), B.1.351 S and B1.1.7 S pseudotypes VSV viruses were prepared as described previously ^54,86^. Briefly, 293T cells in DMEM supplemented with 10% FBS, 1% PenStrep seeded in 10-cm dishes were transfected with the plasmid encoding for the corresponding S glycoprotein using lipofectamine 2000 (Life Technologies) following manufacturer’s indications. One day post-transfection, cells were infected with VSV(G*ΔG-luciferase) and after 2 h were washed five times with DMEM before adding medium supplemented with anti-VSV-G antibody (I1-mouse hybridoma supernatant, CRL-2700, ATCC). Virus pseudotypes were harvested 18-24 h post-inoculation, clarified by centrifugation at 2,500 x g for 5 min, filtered through a 0.45 μm cut off membrane, concentrated 10 times with a 30 kDa cut off membrane, aliquoted and stored at −80°C.

### Pseudovirus Neutralization

HEK-hACE2 cells were cultured in DMEM with 10% FBS (Hyclone) and 1% PenStrep with 8% CO_2_ in a 37°C incubator (ThermoFisher). One day prior to infection, 40 μL of poly-lysine (Sigma) was placed into 96-well plates and incubated with rotation for 5 min. Poly-lysine was removed, plates were dried for 5 min then washed 1 × with water prior to plating cells. The following day, cells were checked to be at 80% confluence. In a half-area 96-well plate a 1:3 serial dilution of sera was made in DMEM in 22 μL final volume. 22 μL of diluted pseudovirus was then added to the serial dilution and incubated at room temperature for 30-60 min. HEK-hACE2 plate media was removed and 40 μL of the sera/virus mixture was added to the cells and incubated for 2 h at 37°C with 8% CO_2_. Following incubation, 40 μL 20% FBS and 2% PenStrep containing DMEM was added to the cells for 48 h. Following the 48-h infection, One-Glo-EX (Promega) was added to the cells in half culturing volume (40 μL added) and incubated in the dark for 5 min prior to reading on a BioTek plate reader.

For VSV-pseudotypes neutralization, HEK-hACE2 in DMEM supplemented with 10% FBS, 1% PenStrep were seeded at 40,000 cells/well into clear bottom white walled 96-well plates and cultured overnight at 37°C. Twelve-point 3-fold serial dilutions of the corresponding mAb were prepared in DMEM and pseudotyped VSV viruses were added 1:1 to each mAb dilution in the presence of anti-VSV-G antibody from I1-mouse hybridoma supernatant diluted 25 times (total volume: 50 µl). After 45 min incubation at 37 °C, 40 µl of the mixture were added to the cells and 2 h post-infection, 40 μL DMEM was added. After 17-20 h 40 μL/well of One-Glo-EX substrate (Promega) was added to the cells and incubated in the dark for 5-10 min prior reading on a BioTek plate reader.

Measurements were done in at least duplicate. Relative luciferase units were plotted and normalized in Prism (GraphPad) using as zero value cells alone or infected with supernatants of non-tranfected cells infected with VSV(G*ΔG-luciferase) VSV(G*ΔG-luciferase) in the presence of anti-VSV-G antibody and as 100% value cells infected with virus alone. Nonlinear regression of log(inhibitor) versus normalized response was used to determine IC_50_ values from curve fits. Kruskal Wallis tests were used to compare two groups to determine whether they were statistically different.

### Sarbecovirus biolayer interferometry binding analysis

Purification of Fabs from NHP serum was adapted from (Boyoglu-Barnum et al., 2020). Briefly, 1 mL of day 70 serum was diluted to 10 mL with PBS and incubated with 1 mL of 3× PBS-washed protein A beads (GenScript) with agitation overnight at 37°C. The next day beads were thoroughly washed with PBS using a gravity flow column and bound Abs were eluted with 0.1 M glycine pH 3.5 into 1M Tris-HCl (pH 8.0) to a final concentration of 100 mM. Serum and early washes that flowed through were re-bound to beads overnight again for a second, repeat elution. IgGs were concentrated (Amicon 30 kDa) and buffer exchanged into PBS. 2× digestion buffer (40 mM sodium phosphate pH 6.5, 20 mM EDTA, 40 mM cysteine) was added to concentrated and pooled IgGs. 500 μL of resuspended immobilized papain resin (ThermoFisher Scientific) freshly washed in 1× digestion buffer (20 mM sodium phosphate, 10 mM EDTA, 20 mM cysteine, pH 6.5) was added to purified IgGs in 2× digestion buffer and samples were agitated for 5 h at 37°C. The supernatant was separated from resin and resin washes were collected and pooled with the resin flow through. Pooled supernatants were sterile-filtered at 0.22 μm and applied 6× to PBS-washed protein A beads in a gravity flow column. The column was eluted as described above and the papain procedure repeated overnight with undigested IgGs to increase yield. The protein A flowthroughs were pooled, concentrated (using an Amicon 10 kDa), and buffer exchanged into PBS. Purity was checked by SDS-PAGE.

Assays were performed and analyzed using biolayer interferometry on an Octet Red 96 System (Pall Forte Bio/Sartorius) at ambient temperature with shaking at 1000 rpm. Different Sarbeco RBDs were diluted with different acetate buffers and were applied to a black 96-well Greiner Bio-one microplate at 200ul per well. GD-Pangolin RBD was diluted in ph6 buffer to 5ug/ml, RmNY02 were diluted in ph5 to 25ug/ml, WIV16 was diluted in ph5 to 10ug/ml, SARS-CoV-2 was diluted in ph6 to 5ug/ml, RaTG13 was diluted in ph6 to 10ug/ml, RaTG13 was diluted in ph6 to 10ug/ml, SARS-Cov was diluted in ph6 to 50ug/ml, and finally ZXC21 was diluted in ph6 to 10ug/ml. AR2G biosensors (ForteBio/Sartorius) following 600 second hydration were normalized in water for 180 seconds. Then tips were NHS-EDC activated for 300 seconds and the different Sarbecovirus RBDs were loaded up to a 1.50nm threshold for up to 600 seconds. Immobilized RBDs on the tips were quenched for 300 seconds in ethanolamine and dipped into kinetics buffer for a 60 second baseline. The association step was performed by dipping the mobilized RBDs into diluted purified polyclonal pigtail macaque IgGs for 600 seconds. Dissociation was measured by inserting the biosensors in kinetics buffer for 600 seconds. The data were baseline subtracted and the plots fitted using the Pall ForteBio/Sartorius analysis software (version 12.0).

### Cocktail and mosaic bio-layer interferometry (antigenicity)

Binding of hACE2-Fc to monovalent RBD-I53-50 nanoparticles, mosaic-RBD-I53-50 nanoparticles, and cocktail of RBD-nanoparticles was analyzed for antigenicity experiments and real-time stability studies using an Octet Red 96 System (Pall FortéBio/Sartorius) at ambient temperature with shaking at 1000 rpm. Protein samples were diluted to 100 nM in Kinetics buffer (1× HEPES-EP+ (Pall Forté Bio), 0.05% nonfat milk, and 0.02% sodium azide). Buffer, receptor, and analyte were then applied to a black 96-well Greiner Bio-one microplate at 200 µL per well. Protein A biosensors (FortéBio/Sartorius) were first hydrated for 10 minutes in Kinetics buffer, then dipped into hACE2-Fc diluted to 10 µg/mL in Kinetics buffer in the immobilization step. After 150 seconds, the tips were transferred to Kinetics buffer for 60 seconds to reach a baseline. The association step was performed by dipping the loaded biosensors into the immunogens for 300 seconds, and subsequent dissociation was performed by dipping the biosensors back into Kinetics buffer for an additional 300 seconds. The data were baseline subtracted prior for plotting using the FortéBio analysis software (version 12.0). Plots in **Extended Data Fig. 4** show the 600 seconds of association and dissociation.

### Sandwich bio-layer interferometry (mosaic display antigenicity)

Binding of hACE2-Fc or S2H14 mAb, and S230 Fab to WIV1-RBD-I53-50, RaTG13-RBD-I53-50, SARS-CoV-SARS-CoV2-RBD-I5350, SARS-CoV2-I5350, and mosaic-RBD-I53-50 nanoparticles were analyzed for co-display of RBDs using an Octet Red 96 System (Pall FortéBio/Sartorius) at ambient temperature with shaking at 1000 rpm. Nanoparticles were diluted to 100 nM in Kinetics buffer. Kinetics buffer, mAb, nanoparticles and Fab were then applied to a black 96-well Greiner Bio-one microplate at 200 µL per well. Protein A biosensors (FortéBio/Sartorius) were first hydrated for 10 minutes in Kinetics buffer, then dipped into hACE2-Fc or S2H14 mAb diluted to 10 µg/mL in Kinetics buffer in the immobilization step. After 150 seconds, the tips were transferred to Kinetics buffer for 60 seconds to reach a baseline. The receptor or mAb was then loaded with nanoparticle by dipping the loaded biosensors into the immunogens for 300 seconds, and subsequent baseline was performed by dipping the biosensors back into the Kinetics buffer for an additional 60 seconds. Association of S230 Fab diluted to 100 nM in Kinetics buffer was then measured for 300 seconds and subsequent dissociation in Kinetics buffer of S230 Fab for 300 seconds. The data were baseline subtracted prior for plotting using the FortéBio analysis software (version 12.0). Plots in Extended Data Fig. 6 exclude the initial mAb loading and the first baseline.

### Cocktail and mosaic negative stain electron microscopy

Monovalent RBD-I53-50 nanoparticles, mosaic-RBD-I53-50 nanoparticles, and cocktail of RBD-nanoparticles were first diluted to 75 µg/mL in 50 mM Tris pH 7.4, 185 mM NaCl, 100 mM Arginine, 4.5% v/v Glycerol, 0.75% w/v CHAPS prior to application of 3 µL of sample onto freshly glow-discharged 300-mesh copper grids. Sample was incubated on the grid for 1 minute before the grid was dipped in a 50 µL droplet of water and excess liquid blotted away with filter paper (Whatman). The grids were then dipped into 6 µL of 0.75% w/v uranyl formate stain. Stain was blotted off with filter paper, then the grids were dipped into another 6 µL of stain and incubated for ∼70 seconds. Finally, the stain was blotted away and the grids were allowed to dry for 1 minute. Prepared grids were imaged in a Talos model L120C electron microscope at 57,000× (nanoparticles).

### Cocktail and mosaic dynamic light scattering

Dynamic Light Scattering (DLS) was used to measure hydrodynamic diameter (Dh) and % Polydispersity (%Pd) of monovalent RBD-I53-50 nanoparticles, mosaic-RBD-I53-50 nanoparticles, and cocktail of RBD-nanoparticles on an UNcle Nano-DSF (UNchained Laboratories). Sample was applied to a 8.8 µL quartz capillary cassette (UNi, UNchained Laboratories) and measured with 10 acquisitions of 5 seconds each, using auto-attenuation of the laser. Increased viscosity due to 4.5% v/v glycerol in the RBD nanoparticle buffer was accounted for by the UNcle Client software in Dh measurements.

### Mouse immunizations and challenge

At six weeks of age, 8 female BALB/c mice per dosing group were vaccinated with a prime immunization, and three weeks later mice were boosted with a second vaccination (IACUC protocol 4470.01). Prior to inoculation, immunogen suspensions were gently mixed 1:1 vol/vol with AddaVax adjuvant (Invivogen, San Diego, CA) to reach a final concentration of 0.01 mg/mL antigen. Mice were injected intramuscularly into the gastrocnemius muscle of each hind leg using a 27-gauge needle (BD, San Diego, CA) with 50 μL per injection site (100 μL total) of immunogen under isoflurane anesthesia. To obtain sera all mice were bled two weeks after prime and boost immunizations. Blood was collected via submental venous puncture and rested in 1.5 mL plastic Eppendorf tubes at room temperature for 30 min to allow for coagulation. Serum was separated from hematocrit via centrifugation at 2,000 g for 10 min. Complement factors and pathogens in isolated serum were heat-inactivated via incubation at 56°C for 60 min. Serum was stored at 4°C or −80°C until use. The study was repeated twice. Five weeks post-boost, mice were exported from Comparative Medicine Facility at the University of Washington, Seattle, WA to an AAALAC accredited Animal Biosafety Level 3 (ABSL3) Laboratory at the University of North Carolina, Chapel Hill. After a 7-day acclimation time, mice were anesthetized with a mixture of ketamine/xylazine and challenged intranasally with 10^5^ plaque-forming units (pfu) of mouse-adapted SARS-CoV-2 MA10 or SARS-CoV MA15 strain for the evaluation of vaccine efficacy (IACUC protocol 20-114.0). After infection, body weight and congestion score were monitored daily until the termination of the study two days post-infection, when lung and nasal turbinate tissues were harvested to evaluate the viral load by plaque assay.

## Quantification and Statistical Analysis

Statistical details of experiments can be found in the figure legends. For NHP experiments, 2 or 5 sera samples were used and experiments were done in duplicate. For mouse ELISAs and neutralization experiments, sera from 8 BALB/c animals were used and experiments were completed in at least duplicate. Geometric mean titers were calculated. Mann-Whitney or Kruskal Wallis tests were performed to compare two groups to determine whether they were statistically different for ELISA and neutralization experiments. Significance is indicated with stars: ^∗^, p < 0.05; ^∗∗∗∗^ p < 0.0001.

## References

Zhou, P. et al. A pneumonia outbreak associated with a new coronavirus of probable bat origin. Nature (2020) doi:10.1038/s41586-020-2012-7.

Zhou, H. et al. A Novel Bat Coronavirus Closely Related to SARS-CoV-2 Contains Natural Insertions at the S1/S2 Cleavage Site of the Spike Protein. Curr. Biol. 30, 2196–2203.e3 (2020).

Menachery, V. D. et al. SARS-like WIV1-CoV poised for human emergence. Proc. Natl. Acad. Sci. U. S. A. 113, 3048–3053 (2016).

Menachery, V. D. et al. A SARS-like cluster of circulating bat coronaviruses shows potential for human emergence. Nat. Med. 21, 1508–1513 (2015).

Walls, A. C. et al. Structure, Function, and Antigenicity of the SARS-CoV-2 Spike Glycoprotein. Cell 181, 281–292.e6 (2020).

Wrapp, D. et al. Cryo-EM structure of the 2019-nCoV spike in the prefusion conformation. Science 367, 1260–1263 (2020).

Letko, M., Marzi, A. & Munster, V. Functional assessment of cell entry and receptor usage for SARS-CoV-2 and other lineage B betacoronaviruses. Nature Microbiology (2020) doi:10.1038/s41564-020-0688-y.

Hoffmann, M. et al. SARS-CoV-2 Cell Entry Depends on ACE2 and TMPRSS2 and Is Blocked by a Clinically Proven Protease Inhibitor. Cell 181, 271–280.e8 (2020).

Wang, Q. et al. Structural and Functional Basis of SARS-CoV-2 Entry by Using Human ACE2. Cell 181, 894–904.e9 (2020).

Yan, R. et al. Structural basis for the recognition of SARS-CoV-2 by full-length human ACE2. Science 367, 1444–1448 (2020).

Shang, J. et al. Structural basis of receptor recognition by SARS-CoV-2. Nature (2020) doi:10.1038/s41586-020-2179-y.

Lan, J. et al. Structure of the SARS-CoV-2 spike receptor-binding domain bound to the ACE2 receptor. Nature (2020) doi:10.1038/s41586-020-2180-5.

Piccoli, L. et al. Mapping Neutralizing and Immunodominant Sites on the SARS-CoV-2 Spike Receptor-Binding Domain by Structure-Guided High-Resolution Serology. Cell 183, 1024–1042.e21 (2020).

Greaney, A. J. et al. Comprehensive mapping of mutations to the SARS-CoV-2 receptor-binding domain that affect recognition by polyclonal human serum antibodies. Cold Spring Harbor Laboratory 2020.12.31.425021 (2021) doi:10.1101/2020.12.31.425021.

Tortorici, M. A. et al. Ultrapotent human antibodies protect against SARS-CoV-2 challenge via multiple mechanisms. Science 370, 950–957 (2020).

Brouwer, P. J. M. et al. Potent neutralizing antibodies from COVID-19 patients define multiple targets of vulnerability. Science (2020) doi:10.1126/science.abc5902.

Zost, S. J. et al. Potently neutralizing and protective human antibodies against SARS-CoV-2. Nature 584, 443–449 (2020).

Pinto, D. et al. Cross-neutralization of SARS-CoV-2 by a human monoclonal SARS-CoV antibody. Nature 583, 290–295 (2020).

Wec, A. Z. et al. Broad neutralization of SARS-related viruses by human monoclonal antibodies. Science (2020) doi:10.1126/science.abc7424.

Rogers, T. F. et al. Isolation of potent SARS-CoV-2 neutralizing antibodies and protection from disease in a small animal model. Science (2020) doi:10.1126/science.abc7520.

Barnes, C. O. et al. SARS-CoV-2 neutralizing antibody structures inform therapeutic strategies. Nature 588, 682–687 (2020).

Baum, A. et al. REGN-COV2 antibodies prevent and treat SARS-CoV-2 infection in rhesus macaques and hamsters. Science (2020) doi:10.1126/science.abe2402.

Corbett, K. S. et al. SARS-CoV-2 mRNA vaccine design enabled by prototype pathogen preparedness. Nature 586, 567–571 (2020).

Corbett, K. S. et al. Evaluation of the mRNA-1273 Vaccine against SARS-CoV-2 in Nonhuman Primates. N. Engl. J. Med. 383, 1544–1555 (2020).

Jackson, L. A. et al. An mRNA Vaccine against SARS-CoV-2 - Preliminary Report. N. Engl. J. Med. (2020) doi:10.1056/NEJMoa2022483.

Polack, F. P. et al. Safety and Efficacy of the BNT162b2 mRNA Covid-19 Vaccine. N. Engl. J. Med. 383, 2603–2615 (2020).

Mercado, N. B. et al. Single-shot Ad26 vaccine protects against SARS-CoV-2 in rhesus macaques. Nature 586, 583–588 (2020).

Yu, J. et al. DNA vaccine protection against SARS-CoV-2 in rhesus macaques. Science (2020) doi:10.1126/science.abc6284.

Tostanoski, L. H. et al. Ad26 vaccine protects against SARS-CoV-2 severe clinical disease in hamsters. Nat. Med. 26, 1694–1700 (2020).

Hou, Y. J. et al. SARS-CoV-2 D614G variant exhibits efficient replication ex vivo and transmission in vivo. Science 370, 1464–1468 (2020).

Korber, B. et al. Tracking Changes in SARS-CoV-2 Spike: Evidence that D614G Increases Infectivity of the COVID-19 Virus. Cell 182, 812–827.e19 (2020).

Plante, J. A. et al. Spike mutation D614G alters SARS-CoV-2 fitness. Nature (2020) doi:10.1038/s41586-020-2895-3.

Yurkovetskiy, L. et al. Structural and Functional Analysis of the D614G SARS-CoV-2 Spike Protein Variant. Cell 183, 739–751.e8 (2020).

Baum, A. et al. Antibody cocktail to SARS-CoV-2 spike protein prevents rapid mutational escape seen with individual antibodies. Science (2020) doi:10.1126/science.abd0831.

Weisblum, Y. et al. Escape from neutralizing antibodies by SARS-CoV-2 spike protein variants. Elife 9, e61312 (2020).

Li, Q. et al. The Impact of Mutations in SARS-CoV-2 Spike on Viral Infectivity and Antigenicity. Cell 182, 1284–1294.e9 (2020).

Collier, D. A. et al. SARS-CoV-2 B.1.1.7 escape from mRNA vaccine-elicited neutralizing antibodies. medRxiv 2021.01.19.21249840 (2021).

Greaney, A. J. et al. Complete Mapping of Mutations to the SARS-CoV-2 Spike Receptor-Binding Domain that Escape Antibody Recognition. Cell Host Microbe (2020) doi:10.1016/j.chom.2020.11.007.

Dong, J. et al. Genetic and structural basis for recognition of SARS-CoV-2 spike protein by a two-antibody cocktail. bioRxiv (2021) doi:10.1101/2021.01.27.428529.

Tegally, H. et al. Emergence and rapid spread of a new severe acute respiratory syndrome-related coronavirus 2 (SARS-CoV-2) lineage with multiple spike mutations in South Africa. medRxiv 2020.12.21.20248640 (2020).

Davies, N. G. et al. Estimated transmissibility and severity of novel SARS-CoV-2 Variant of Concern 202012/01 in England. medRxiv 2020.12.24.20248822 (2020).

Faria, N. R. et al. Genomic characterisation of an emergent SARS-CoV-2 lineage in Manaus: preliminary findings. January 12, 2021 (2021).

Wibmer, C. K. et al. SARS-CoV-2 501Y.V2 escapes neutralization by South African COVID-19 donor plasma. bioRxiv 2021.01.18.427166 (2021).

Shapiro, L., Sheng, Z., Nair, M. S., Huang, Y. & Ho, D. D. Increased resistance of SARS-CoV-2 variants B. 1.351 and B. 1.1. 7 to antibody neutralization. bioRxiv (2021).

Wang, Z. et al. mRNA vaccine-elicited antibodies to SARS-CoV-2 and circulating variants. bioRxiv (2021) doi:10.1101/2021.01.15.426911.

Collier, D. A. et al. Sensitivity of SARS-CoV-2 B.1.1.7 to mRNA vaccine-elicited antibodies. Nature (2021) doi:10.1038/s41586-021-03412-7.

Walls, A. C. et al. Elicitation of potent neutralizing antibody responses by designed protein nanoparticle vaccines for SARS-CoV-2. bioRxiv 2020.08.11.247395 (2020).

Bale, J. B. et al. Accurate design of megadalton-scale two-component icosahedral protein complexes. Science 353, 389–394 (2016).

Dinnon, K. H. et al. A mouse-adapted model of SARS-CoV-2 to test COVID-19 countermeasures. Nature (2020) doi:10.1038/s41586-020-2708-8.

S Arunachalam, P. et al. Adjuvanting a subunit SARS-CoV-2 nanoparticle vaccine to induce protective immunity in non-human primates. bioRxiv (2021) doi:10.1101/2021.02.10.430696.

Tortorici, M. A. & Veesler, D. Structural insights into coronavirus entry. Adv. Virus Res. 105, 93–116 (2019).

Walls, A. C. et al. Cryo-electron microscopy structure of a coronavirus spike glycoprotein trimer. Nature 531, 114–117 (2016).

Rappazzo, C. G. et al. An Engineered Antibody with Broad Protective Efficacy in Murine Models of SARS and COVID-19. bioRxiv (2020) doi:10.1101/2020.11.17.385500.

McCallum, M. et al. N-terminal domain antigenic mapping reveals a site of vulnerability for SARS-CoV-2. bioRxiv 2021.01.14.426475 (2021).

McCarthy, K. R. et al. Recurrent deletions in the SARS-CoV-2 spike glycoprotein drive antibody escape. Science (2021) doi:10.1126/science.abf6950.

Andreano, E. et al. SARS-CoV-2 escape in vitro from a highly neutralizing COVID-19 convalescent plasma. bioRxiv 2020.12.28.424451 (2020).

Avanzato, V. A. et al. Case Study: Prolonged infectious SARS-CoV-2 shedding from an asymptomatic immunocompromised cancer patient. Cell (2020) doi:10.1016/j.cell.2020.10.049.

Choi, B. et al. Persistence and Evolution of SARS-CoV-2 in an Immunocompromised Host. N. Engl. J. Med. (2020) doi:10.1056/NEJMc2031364.

Ho, D. et al. Increased Resistance of SARS-CoV-2 Variants B.1.351 and B.1.1.7 to Antibody Neutralization. doi:10.21203/rs.3.rs-155394/v1.

Millet, J. K. & Whittaker, G. R. Murine Leukemia Virus (MLV)-based Coronavirus Spike-pseudotyped Particle Production and Infection. Bio Protoc 6, (2016).

Leist, S. R. et al. A Mouse-Adapted SARS-CoV-2 Induces Acute Lung Injury and Mortality in Standard Laboratory Mice. Cell 183, 1070–1085.e12 (2020).

Starr, T. N. et al. Deep Mutational Scanning of SARS-CoV-2 Receptor Binding Domain Reveals Constraints on Folding and ACE2 Binding. Cell 182, 1295–1310.e20 (2020).

Hsieh, C. L. et al. Structure-based design of prefusion-stabilized SARS-CoV-2 spikes. Science (2020) doi:10.1126/science.abd0826.

Greaney, A. J. et al. Comprehensive mapping of mutations in the SARS-CoV-2 receptor-binding domain that affect recognition by polyclonal human plasma antibodies. Cell Host Microbe (2021) doi:10.1016/j.chom.2021.02.003.

Wang, Z. et al. mRNA vaccine-elicited antibodies to SARS-CoV-2 and circulating variants. Nature (2021) doi:10.1038/s41586-021-03324-6.

Thomson, E. C. et al. Circulating SARS-CoV-2 spike N439K variants maintain fitness while evading antibody-mediated immunity. Cell (2021) doi:10.1016/j.cell.2021.01.037.

Liu, Z. et al. Identification of SARS-CoV-2 spike mutations that attenuate monoclonal and serum antibody neutralization. Cell Host Microbe (2021) doi:10.1016/j.chom.2021.01.014.

Pallesen, J. et al. Immunogenicity and structures of a rationally designed prefusion MERS-CoV spike antigen. Proc. Natl. Acad. Sci. U. S. A. 114, E7348–E7357 (2017).

Stamatatos, L. et al. Antibodies elicited by SARS-CoV-2 infection and boosted by vaccination neutralize an emerging variant and SARS-CoV-1. medRxiv (2021).

Jabal, K. A. et al. Impact of age, ethnicity, sex and prior infection status on immunogenicity following a single dose of the BNT162b2 mRNA COVID-19 vaccine: real-world evidence from healthcare workers, Israel, December 2020 to January 2021. Eurosurveillance vol. 26 (2021).

Krammer, F., Srivastava, K., Simon, V. & Others. Robust spike antibody responses and increased reactogenicity in seropositive individuals after a single dose of SARS-CoV-2 mRNA vaccine. medRxiv (2021).

Lam, T. T. et al. Identifying SARS-CoV-2 related coronaviruses in Malayan pangolins. Nature (2020) doi:10.1038/s41586-020-2169-0.

Boyoglu-Barnum, S. et al. Elicitation of broadly protective immunity to influenza by multivalent hemagglutinin nanoparticle vaccines. bioRxiv 2020.05.30.125179 (2020).

Kanekiyo, M. et al. Mosaic nanoparticle display of diverse influenza virus hemagglutinins elicits broad B cell responses. Nat. Immunol. 20, 362–372 (2019).

Cohen, A. A. et al. Construction, characterization, and immunization of nanoparticles that display a diverse array of influenza HA trimers. doi:10.1101/2020.01.18.911388.

Cohen, A. A. et al. Mosaic nanoparticles elicit cross-reactive immune responses to zoonotic coronaviruses in mice. Science 371, 735–741 (2021).

Walls, A. C. et al. Unexpected Receptor Functional Mimicry Elucidates Activation of Coronavirus Fusion. Cell 176, 1026–1039.e15 (2019).

Traggiai, E. et al. An efficient method to make human monoclonal antibodies from memory B cells: potent neutralization of SARS coronavirus. Nat. Med. 10, 871–875 (2004).

Rockx, B. et al. Structural basis for potent cross-neutralizing human monoclonal antibody protection against lethal human and zoonotic severe acute respiratory syndrome coronavirus challenge. J. Virol. 82, 3220–3235 (2008).

Roberts, A. et al. A mouse-adapted SARS-coronavirus causes disease and mortality in BALB/c mice. PLoS Pathog. 3, e5 (2007).

Wu, K. et al. Serum Neutralizing Activity Elicited by mRNA-1273 Vaccine - Preliminary Report. N. Engl. J. Med. (2021) doi:10.1056/NEJMc2102179.

Dejnirattisai, W. et al. The antigenic anatomy of SARS-CoV-2 receptor binding domain. Cell (2021) doi:10.1016/j.cell.2021.02.032.

Walls, A. C. et al. Elicitation of Potent Neutralizing Antibody Responses by Designed Protein Nanoparticle Vaccines for SARS-CoV-2. Cell 183, 1367–1382.e17 (2020).

Starr, T. N. et al. Prospective mapping of viral mutations that escape antibodies used to treat COVID-19. bioRxiv (2020) doi:10.1101/2020.11.30.405472.

Otwinowski, J., McCandlish, D. M. & Plotkin, J. B. Inferring the shape of global epistasis. Proc. Natl. Acad. Sci. U. S. A. 115, E7550–E7558 (2018).

Sauer, M. M. et al. Structural basis for broad coronavirus neutralization. bioRxiv (2020) doi:10.1101/2020.12.29.424482.

